# *Toxoplasma gondii* ROP1 subverts murine and human innate immune restriction

**DOI:** 10.1101/2022.03.21.485090

**Authors:** Simon Butterworth, Francesca Torelli, Eloise J. Lockyer, Jeanette Wagener, Ok-Ryul Song, Malgorzata Broncel, Matt R. G. Russell, Joanna C. Young, Moritz Treeck

## Abstract

*Toxoplasma gondii* is an intracellular parasite that can infect many different host species and is a cause of significant human morbidity worldwide. *T. gondii* secretes a diverse array of effector proteins into the host cell which are critical for infection; however, the vast majority of these secreted proteins are uncharacterised. Here, we carried out a pooled CRISPR knockout screen in the *T. gondii* Prugniaud strain *in vivo* to identify secreted proteins that contribute to parasite immune evasion in the host. We identify 22 putative virulence factors and demonstrate that ROP1, the first-identified rhoptry protein of *T. gondii*, has a previously unrecognised role in parasite resistance to interferon gamma-mediated innate immune restriction. This function is conserved in the highly virulent RH strain of *T. gondii* and contributes to parasite growth in both murine and human macrophages. While ROP1 affects the morphology of rhoptries, from where the protein is secreted, it does not affect rhoptry secretion. ROP1 interacts with the host cell protein C1QBP, which appears to facilitate parasite immune evasion. In summary, we identify 22 secreted proteins which contribute to parasite growth *in vivo* and show that ROP1 is an important and previously overlooked effector in counteracting both murine and human innate immunity.

## INTRODUCTION

*Toxoplasma gondii* is a single-celled intracellular parasite which is remarkable in its ability to infect any warm-blooded animal, including humans. In intermediate hosts, *T. gondii* tachyzoites must evade host immune clearance long enough to disseminate throughout the host organism and differentiate into the cyst-forming bradyzoites, which can be transmitted to the definitive feline host (Dubey 2014).

To this end, *T. gondii* secretes effector proteins into the host cell which modulate and counteract host innate immunity pathways (Lima and Lodoen 2019; Frickel and Hunter 2021). These effector proteins are secreted from the rhoptries and dense granules, specialised secretory organelles found in the *Apicomplexa*.

Several effectors have previously been identified by quantitative trait locus (QTL) mapping following genetic crosses between strains of *T. gondii* with differing virulence in mouse models of infection (Saeij et al. 2006; Taylor et al. 2006; Behnke et al. 2011). This approach led to the discovery of ROP5 and ROP18, rhoptry proteins that cooperate to inhibit loading of host immune-related GTPases (IRGs) onto the parasitophorous vacuole membrane (PVM), and that are the major determinants of virulence in mice between different strains of *T. gondii* (Fentress et al. 2010; Reese et al. 2011; Fleckenstein et al. 2012; Behnke et al. 2012; Niedelman et al. 2012; Behnke et al. 2015). However, genetic cross approaches are limited in that they cannot identify effector proteins with the same function in both parental strains; for example, the dense granule protein GRA12, which has been shown to be a major virulence factor in both Type I and Type II laboratory strains of *T. gondii* (Fox et al. 2019; J.-L. Wang et al. 2020).

Recently, we and others have used targeted, pooled CRISPR knockout screening to identify *T. gondii* genes which are required for survival and growth in mouse models of infection (Young et al. 2019; Sangaré et al. 2019). By comparison to *in vitro* growth phenotypes, it is possible to identify genes which are only required for parasite growth *in vivo*, and thus may have roles in evasion of the host immune response.

These studies primarily targeted genes encoding proteins localised to the rhoptries and dense granules, as these proteins are secreted into the host cell and have the potential to interact with host proteins. However, 142 proteins that have only recently been localised to the rhoptries and dense granules have yet to be characterised (Barylyuk et al. 2020).

To address this knowledge gap, we here use our previously described platform for customisable pooled CRISPR knockout screening in *T. gondii* (Young et al. 2019) to screen an expanded library of rhoptry and dense granule protein-encoding genes for *in vivo* growth phenotypes in the Type II Prugniaud (PRU) strain of *T. gondii*. We report phenotype scores for 164 genes, of which 75 are putative rhoptry/dense granule protein-encoding genes which have not previously been assessed by *in vivo* screening in the PRU strain, and 61 of which have not been assessed in either PRU or the Type I RH strain (Young et al. 2019; Sangaré et al. 2019). We identify 22 effectors putatively required for immune evasion, of which nine have not previously been studied in the PRU strain. These putative effectors include the prototypical rhoptry protein ROP1, whose function has been unknown to date. We demonstrate that ROP1 protects against IFNγ-mediated restriction in human and murine macrophages in both the PRU and RH strains of *T. gondii*. Finally, we show that ROP1 interacts with C1QBP, a host protein previously implicated in numerous host cell pathways, including defence against infection. Deletion of C1QBP enhances IFNγ-mediated restriction of wild-type parasites, but does not further restrict ΔROP1 parasite growth. This indicates that C1QBP is not a classical restriction factor and that the interaction between ROP1 and C1QBP is beneficial for the parasite in IFNγ-activated immune cells.

## RESULTS

### CRISPR Screen

To screen for *T. gondii* effector proteins required for immune evasion, we cloned 906 protospacer sequences targeting 235 rhoptry and dense granule protein-encoding genes into a Cas9-sgRNA vector (Young et al. 2019) **(Supplementary Data 1)**. We transfected the resulting plasmid pool into the Type II PRUΔHXGPRT strain of *T. gondii*, and selected for integration of the plasmids into the parasite genome for six days in human foreskin fibroblasts (HFFs) using a drug resistance marker. The surviving parasites were used to infect five C57BL/6 mice with 200,000 parasites each by injection into the peritoneum. After five days of infection, parasites were recovered from the mice by peritoneal lavage and expanded in HFFs for one passage **(Figure 1A)**. To quantify the growth of parasite mutants in cell culture, sgRNAs were amplified by PCR from the plasmid pool and from genomic DNA extracted from the parasites after the *in vitro* drug selection. To quantify growth *in vivo*, sgRNAs were amplified from the leftover mouse inoculum and from the five recovered *ex vivo* populations. The sgRNAs from each population were sequenced by Illumina sequencing to determine their relative abundance.

**Figure 1:**
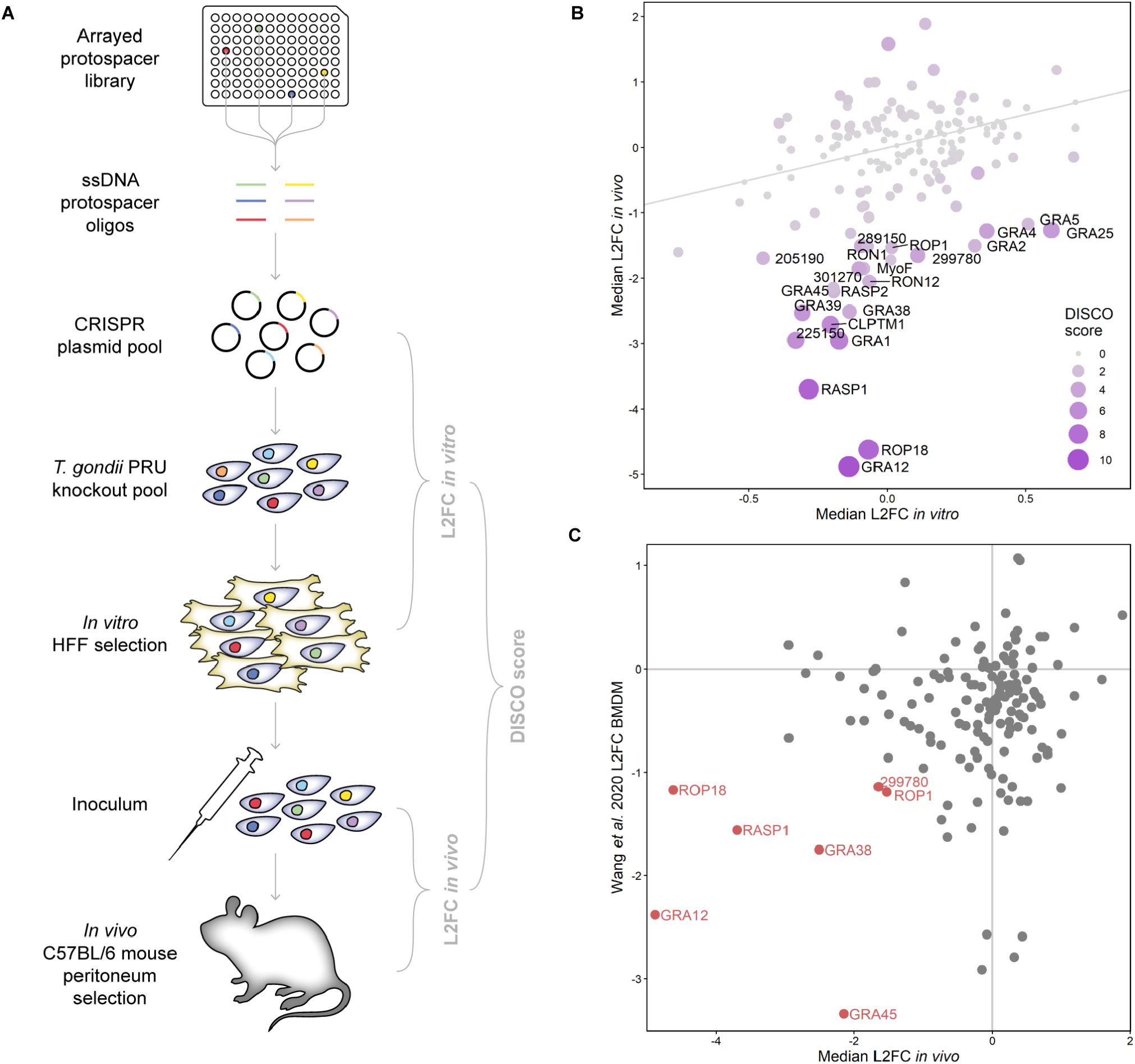
Targeted *in vivo* CRISPR-Cas9 knockout screening of *T. gondii* rhoptry and dense granule proteins. **A.** Schematic of knockout screen workflow. Protospacers encoded on arrayed ssDNA oligonucleotides are assembled by pooled Gibson cloning into a Cas9-sgRNA vector. The resulting plasmid pool is transfected into *T. gondii* PRU and the parasites selected *in vitro* in HFFs for integration for six days. Surviving parasites are used to infect five mice by intraperitoneal injection, recovered after five days and expanded for one further lytic cycle *in vitro*. The sgRNA cassettes are amplified from plasmid or parasite genomic DNA and sequenced to determine the relative abundance of each guide. **B.** Scatter plot of median L2FCs for each gene *in vitro* and *in vivo*. Each point is scaled according to the gene discordance-concordance (DISCO) score. Genes with an *in vivo* L2FC < −1 and DISCO score >2 are labelled. The grey line indicates equal *in vitro* and *in vivo* L2FCs. **C.** Correlation between median L2FCs *in vivo* from this study and L2FCs between IFNγ-stimulated versus unstimulated BMDMs from (Yifan Wang et al. 2020). Genes with a L2FC < −1 in both screens are labelled.

To quantify the contribution of each gene to parasite growth, we calculated a phenotype score as the median log_2_*-*fold-change (L2FC) of the sgRNAs targeting a given gene during the *in vitro* and *in vivo* selections (drug-selected parasites vs. plasmid library and *ex vivo* population vs. inoculum respectively) **(Figure 1B)**. We obtained such scores for 164 genes after filtering to remove genes with fewer than three well-represented sgRNAs **(Supplementary Data 1)**. Both the *in vitro* and *in vivo* phenotype scores correlated strongly with those from our previous study (Young et al. 2019) (Pearson correlation coefficents of 0.70 and 0.88 respectively), indicating that these phenotypes are highly reproducible within this system **(Figure S1A & S1B)**.

To identify genes which contribute to fitness *in vivo* but not *in vitro*, and therefore likely have roles in host immune evasion, we calculated discordance-concordance (DISCO) scores for each gene **(Figure 1B)**. This score takes into account both the *in vitro* and *in vivo* median L2FCs and p-values calculated by paired *t*-test on the log-transformed normalised sgRNA counts. A higher DISCO score indicates more discordant phenotypes, i.e. contributing to fitness in only one condition.

We defined a set of putative immune-evading effectors as the genes with L2FCs *in vivo* less than −1 and a DISCO score greater than 2 **(Supplementary Data 1, annotated in Figure 1B)**. This candidate list contained seven control genes which have previously been shown to be non-essential *in vitro* but contribute to virulence of Type II *T. gondii* strains in mice: GRA12 (Fox et al. 2019), ROP18 (Fox et al. 2016), GRA25 (Shastri et al. 2014), GRA4 (Fox et al. 2019), TGME49_289150 (Young et al. 2019), GRA2 (Fox et al. 2019), and GRA39 (Nadipuram et al. 2016), while excluding 11 genes which have been shown to not affect virulence **(Figure S1C)**. These criteria also identified several genes whose apparent contribution to growth of *T. gondii* PRU *in vivo* has not previously been shown, including RASP1 (Suarez et al. 2019), TGME49_225150, RON12 (Camejo et al. 2014), ROP1 (Ossorio, Schwartzman, and Boothroyd 1992; Saffer et al. 1992; Soldati et al. 1995) and GRA5 (Lecordier et al. 1993).

We were interested in determining whether any of the putative effectors we identified showed a similar phenotype in other strains of *T. gondii*. We initially compared our phenotype scores to a CRISPR knockout screen carried out in the RH strain of *T. gondii* in CD-1 mice; however, the overlap with our targets was only 31 genes (Sangaré et al. 2019). Instead, we compared our data to results from a genome-wide screen carried out in IFNγ-stimulated C57BL/6 bone marrow-derived macrophages (BMDMs), which are thought to recapitulate acute infection in the peritoneum (Yifan Wang et al. 2020) **(Figure 1C)**.

Seven out of 150 unambiguous orthologues present in both screens had L2FCs of less than −1 in both screens, indicating that they are important for immune evasion of both the RH and PRU strains: ROP1, GRA12, ROP18, GRA45, GRA38, RASP1, and TGME9_299780 **(Figure 1C)**. Both GRA12 and ROP18 have been shown to affect virulence in both RH and PRU (J.-L. Wang et al. 2020; Fox et al. 2019; Saeij et al. 2006; Taylor et al. 2006; Young et al. 2019). ROP18 has several functions in the host cell, including in inhibiting loading of host IRGs onto the PVM (Fentress et al. 2010; Fleckenstein et al. 2012). The function of GRA12 in IFNγ resistance is less clear, but appears to depend on the host regulatory IRGs IRGM1/3 (Fox et al. 2019). GRA45, meanwhile, has recently been shown in the RH strain to be required for trafficking of dense granule proteins to the PVM, and likely fulfills the same function in PRU (Yifan Wang et al. 2020).

The functions of GRA38, RASP1, TGME49_299780 and ROP1 are currently unknown, although an interesting rhoptry morphology phenotype has been reported for ROP1 (Soldati et al. 1995). For this reason, we chose to focus the remaining work on ROP1, the first-discovered rhoptry protein of *T. gondii*, which is very highly expressed but whose function has been mysterious. Given the fitness phenotypes described above, we hypothesised that ROP1 contributes to growth of parasites *in vivo* by inhibiting IFNγ-mediated restriction in infected macrophages, potentially through facilitating efficient secretion of rhoptry proteins into the host cell.

### ROP1 localises to the parasitophorous vacuole membrane at 24 hours post-invasion

To investigate the function of ROP1, we generated knockout cell lines in both the PRUΔKU80 and RHΔKU80 strains by replacing the coding sequence of ROP1 with an mCherry-T2A-HXGPRT drug selection cassette. We then complemented these lines with strain-matched ROP1-HA constructs integrated at the UPRT locus. Correct genomic integration of these constructs was verified by PCR, and the expected presence/absence of ROP1 was demonstrated by Western blot and immunofluorescence assay (IFA) **(Figure 2A, Figure S2A & S2B)**.

**Figure 2:**
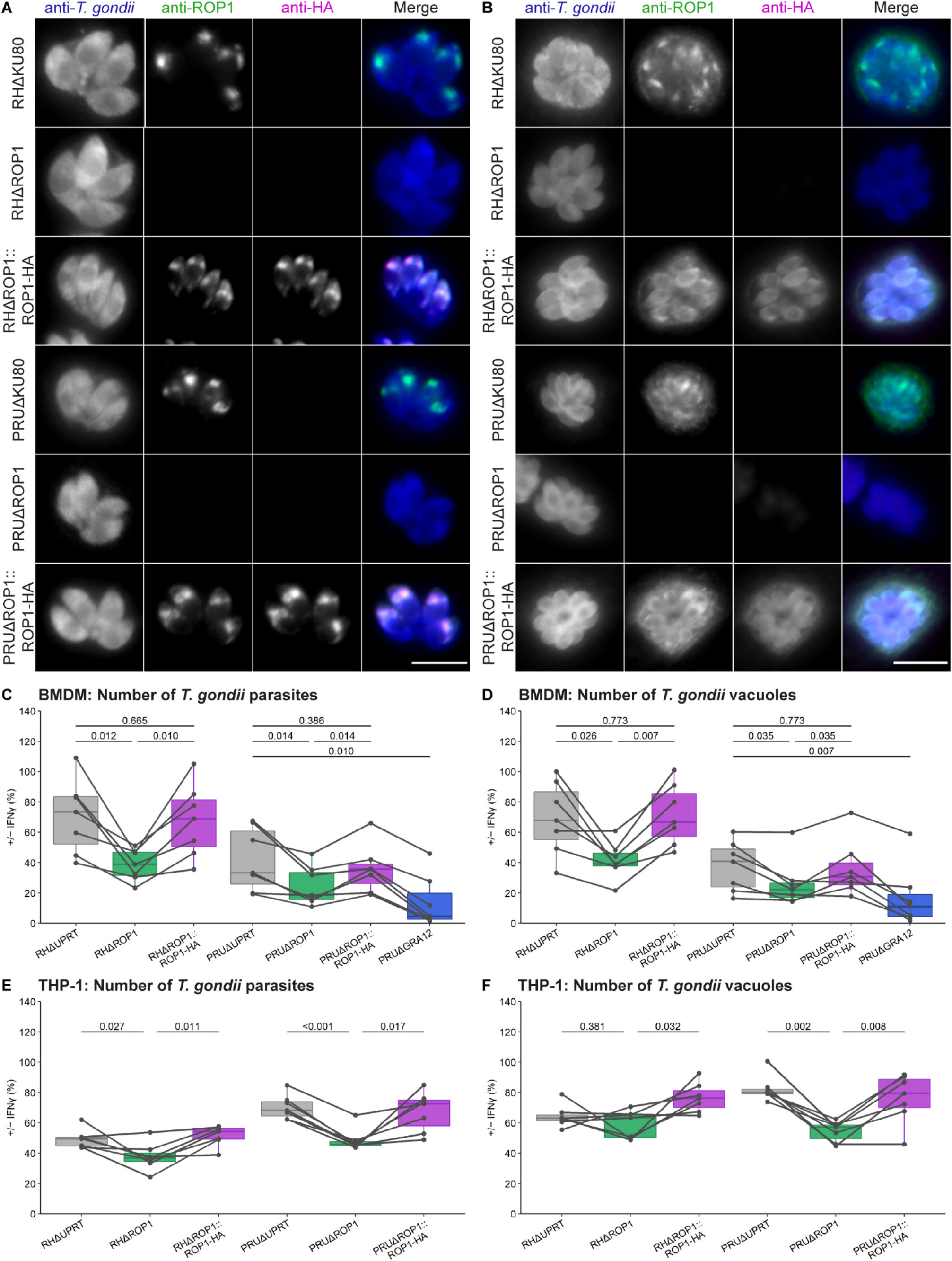
ROP1 contributes to *T. gondii* resistance to IFNγ in murine and human macrophages. **A, B.** Immunofluorescence verification of ROP1 knockout and complemented *T. gondii* cell lines using **A** full permeabilisation (15 minutes 0.2% Triton X-100) or **B** short permeabilisation (1 minute 0.2% Triton X-100). Scale bar = 10 μm. **C, D.** IFNγ-dependent growth restriction of *T. gondii* in BMDMs. BMDMs were stimulated with IFNγ for 24 h, infected with *T. gondii* cell lines for a further 24 h, and parasite growth quantified by automated fluorescence imaging and analysis. Parasite growth in IFNγ-stimulated BMDMs is shown as a percentage of that in unstimulated BMDMs in terms of **C** total parasite number and **D** vacuole number. p-values were calculated by paired two-sided *t-*test with Benjamini-Hochberg adjustment. **E, F.** IFNγ-dependent growth restriction of *T. gondii* in THP-1-derived macrophages. Differentiated THP-1 macrophages were stimulated with IFNγ, infected, and parasite growth quantified as above. Parasite growth in IFNγ-stimulated THP-1 macrophages is shown as a percentage of that in unstimulated macrophages in terms of **E** total parasite number and **F** vacuole number. p-values were calculated by paired two-sided *t-*test with Benjamini-Hochberg adjustment.

We noted that ROP1 is detectable by IFA at the parasitophorous vacuole membrane (PVM) up to at least 24 h post-invasion when the cells are permeabilised for a shorter period (1 min versus 15 min with 0.2% Triton X-100), which allows better visualisation of proteins at the PVM **(Figure 2B)**. This contrasts with a previous report based on immuno-electron microscopy that ROP1 was present on the PVM immediately after invasion but not at 6 h post-invasion (Saffer et al. 1992), but is supported by a recent proximity biotinylation study which demonstrated that ROP1 is accessible to host cytosolic proteins at 24 h post-invasion (Cygan et al., 2021).

Consistent with our CRISPR screen *in vitro* phenotype and a previous study (Soldati et al. 1995), we did not see a major growth defect of the RHΔROP1 or PRUΔROP1 strains compared to the parental strains by plaque assay **(Figure S2C)**, demonstrating that ROP1 is dispensable in the absence of immune pressure.

### ROP1 contributes to *T. gondii* resistance to IFNγ in murine and human macrophages

To test our hypothesis that ROP1 contributes to fitness *in vivo* by protecting the parasite against host IFNγ-mediated restriction, we quantified *T. gondii* growth in primary bone marrow-derived macrophages (BMDMs). BMDMs in 96-well plates were stimulated with IFNγ for 24 h or left unstimulated, then infected with mCherry-expressing parasite strains for a further 24 h. Fluorescence microscopy images of the parasite and host cells were captured using a high-content imaging system and the number of parasites, vacuoles, and host cells determined. The percentage survival of the parasites in IFNγ-stimulated cells was calculated relative to unstimulated cells **(Supplementary Data 2)**.

In terms of total parasite number, survival of both RHΔROP1 and PRUΔROP1 was significantly reduced compared to UPRT knockout controls. RHΔROP1 showed a 32% mean absolute decrease versus RHΔUPRT, and PRUΔROP1 showed a 17% mean absolute decrease versus PRUΔUPRT **(Figure 2C)**. This was rescued by complementation. Consistent with our CRISPR screen, PRUΔROP1 had an intermediate IFNγ resistance phenotype compared to PRUΔGRA12, a virulence factor which has been shown to be highly susceptible to IFNγ-mediated restriction in murine cells (Fox et al. 2019).

The reduced parasite number was the result of reduced vacuole number, but not reduced vacuole size or host cell number **(Figure 2D, Figure S3A & S3B)**, indicating that ROP1 knockout parasites are more susceptible to IFNγ-mediated vacuole destruction rather than growth limitation or host cell death.

We were interested in determining whether ROP1 might contribute to parasite survival in human macrophages, which also restrict parasite growth in an IFNγ-dependent manner but lack the IRG system responsible for this in murine cells (Frickel and Hunter 2021). We stimulated human THP-1-derived macrophages with IFNγ, infected them with mCherry-expressing parasite strains and analysed parasite growth by high-content imaging as above **(Supplementary Data 3)**. As we observed in BMDMs, we found that ΔROP1 parasites of both the RH and PRU strains are more restricted in THP1-derived macrophages than either ΔUPRT or complemented parasite lines **(Figure 2E).** For PRUΔROP1, we observed the same reduction in vacuole number, but not in vacuole size or host cell number, as in BMDMs **(Figure 2F, Figure S3C & S3D)**. For RHΔROP1, we found a significant decrease in the number of vacuoles compared to the complemented line but not compared to RHΔUPRT, and instead observed a modest but significant decrease in vacuole size compared to RHΔUPRT.

Together, these data show that ROP1 contributes to resistance to IFNγ-mediated restriction in both the RH and PRU strains, and in both murine and human macrophages. Enhanced restriction of ΔROP1 parasites is primarily mediated through increased vacuole destruction. In contrast to established rhoptry virulence factors ROP5 and ROP18, the function of ROP1 is likely independent of the IRG genes that act to restrict parasite growth in murine cells, as the IRGs are absent in human macrophages.

### ROP1 affects rhoptry morphology, but not ROP secretion

Knockout of ROP1 in the RH strain has previously been shown to affect the ultrastructure of the rhoptries: wild-type rhoptries have a heterogeneous texture by transmission electron microscopy, whereasin the absence of ROP1 the rhoptries show a homogenously electron-dense structure (Soldati et al. 1995). We were able to reproduce this phenotype previously observed in the RH strain, and additionally show that it is conserved in the Type II PRU strain **(Figure 3A, S4)**.

**Figure 3:**
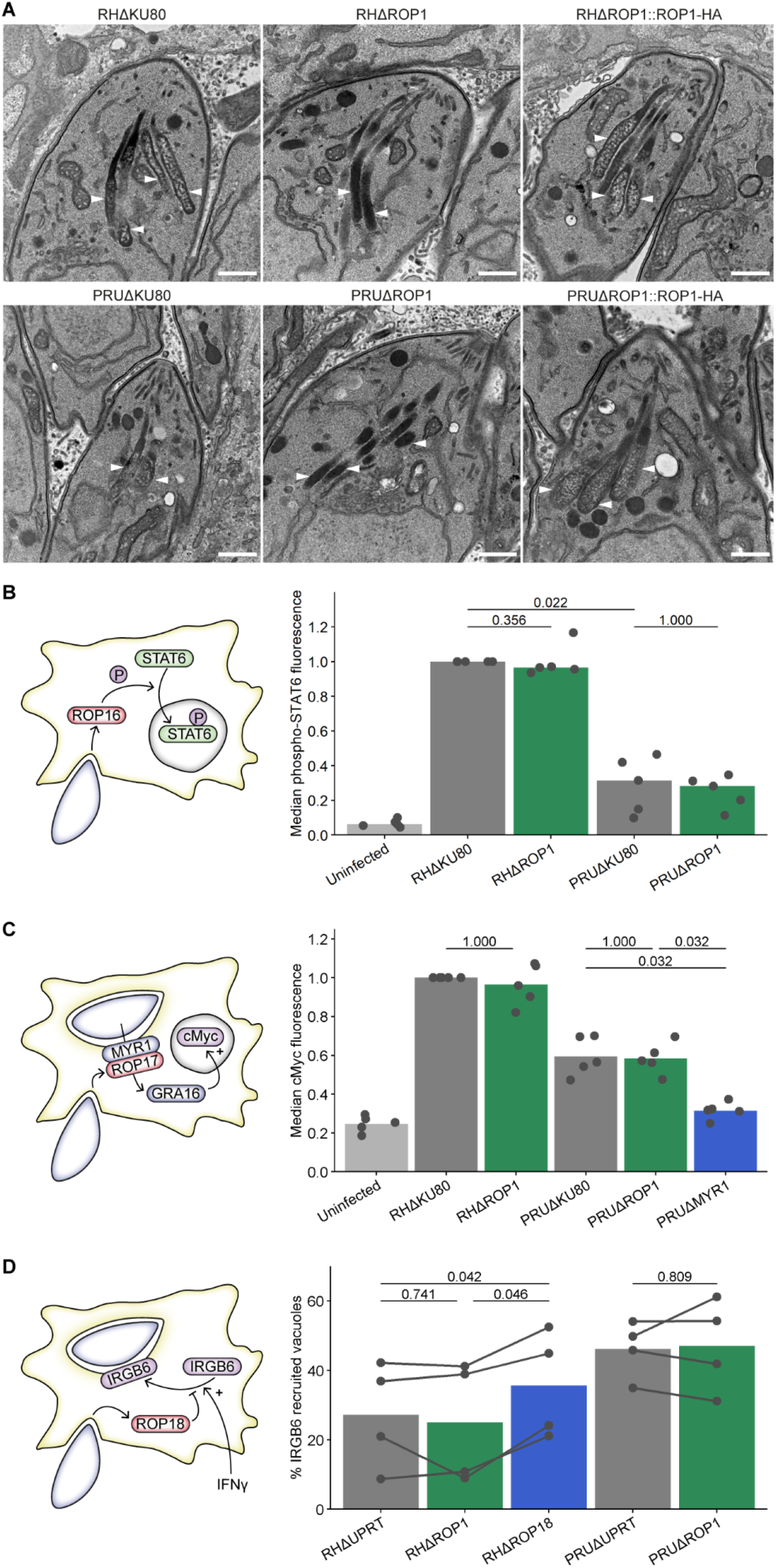
ROP1 knockout alters rhoptry morphology but not ROP secretion. **A.** TEM images of the rhoptries of intracellular tachyzoites. White arrowheads indicate rhoptries. Scale bar = 500 μm. **B.** Normalised median anti-phospho-STAT6 fluorescence intensity of infected HFFs. HFFs were infected for 2 h, fixed with methanol and stained with anti-phospho-STAT6 and anti-*T. gondii*, then analysed by flow cytometry. p-values were calculated by two-sided Wilcoxon rank sum test with Bonferroni correction. **C.** Normalised median nuclear anti-cMyc fluorescence intensity of infected HFFs. HFFs were infected for 24 h in 0.1% FBS medium, fixed and stained with anti-cMyc and anti-*T. gondii*, and the median nuclear anti-cMyc fluorescence intensity was determined from immunofluorescence microscopy images. p-values were calculated by two-sided Wilcoxon rank sum test with Bonferroni correction. **D.** Recruitment of host IRGB6 to *T. gondii* vacuoles in BMDMs. BMDMs were stimulated with IFNγ for 24 h, infected with *T. gondii* cell lines for 1 h, fixed and stained with anti-IRGB6. The percentage of vacuoles decorated with IRGB6 was determined by automated fluorescence imaging and analysis. p-values were calculated by paired two-sided *t-*test with Bonferroni correction.

A subset of rhoptry proteins localised to the apical neck of the rhoptries (RONs) have critical roles in host cell invasion (Boothroyd and Dubremetz 2008). RHΔROP1 parasites have been shown to have the same invasion rate as the parental strain (Soldati et al. 1995), indicating that these parasites have normal secretion of RON proteins. However, given the altered morphology of the rhoptries and rhoptry bulb localisation of ROP1, we hypothesised that secretion of other rhoptry bulb (ROP) proteins might be affected by ROP1 knockout independently of the RON proteins. This would explain the IFNγ-dependent growth defect of ΔROP1 parasites, as other rhoptry bulb proteins have been shown to influence resistance to IFNγ-mediated restriction (e.g. ROP18 (Fentress et al. 2010), ROP54 (Kim et al. 2016)).

As a proxy for secretion, we analysed host cell phenotypes induced by three ROP proteins: phosphorylation of STAT6 by ROP16 (in the RH strain but not PRU) (Saeij et al. 2007), ROP17-dependent induction of cMyc expression in the host nucleus (Panas et al. 2019), and recruitment of IRGB6 to the parasitophorous vacuole membrane, which is inhibited by ROP18 in the RH strain (Fentress et al. 2010) **(Figure 3B, 3C & 3D)**. We did not find a significant difference between ΔROP1 and parental lines in any of these assays, although we observed the expected decrease in host STAT6 phosphorylation for PRUΔKU80 versus RHΔKU80-infected cells, decrease in host cMyc expression for PRUΔMYR1 versus PRUΔKU80-infected cells, and increase in IRGB6 recruitment to RHΔROP18 versus RHΔKU80 vacuoles. It is therefore unlikely that ROP1 has a major function in secretion of ROP proteins. The functional consequences, if any, of the altered rhoptry morphology remain unknown, as neither RON nor ROP secretion is affected. These experiments instead suggest that resistance to IFNγ-mediated restriction is most likely an inherent function of ROP1.

### ROP1 co-immunoprecipitates with host C1QBP

To identify interacting partners of ROP1 that might inform on its function, we generated cell lines in which ROP1 was tagged at the endogenous C-terminus with a single haemagglutinin (HA) epitope. Correct integration of the tagging construct into the genome was confirmed by PCR **(Figure S5A)**, expression of ROP1-HA was demonstrated by Western blot **(Figure S5B)**, and correct localisation of ROP1-HA was determined by IFA **(Figure 4A)**.

**Figure 4:**
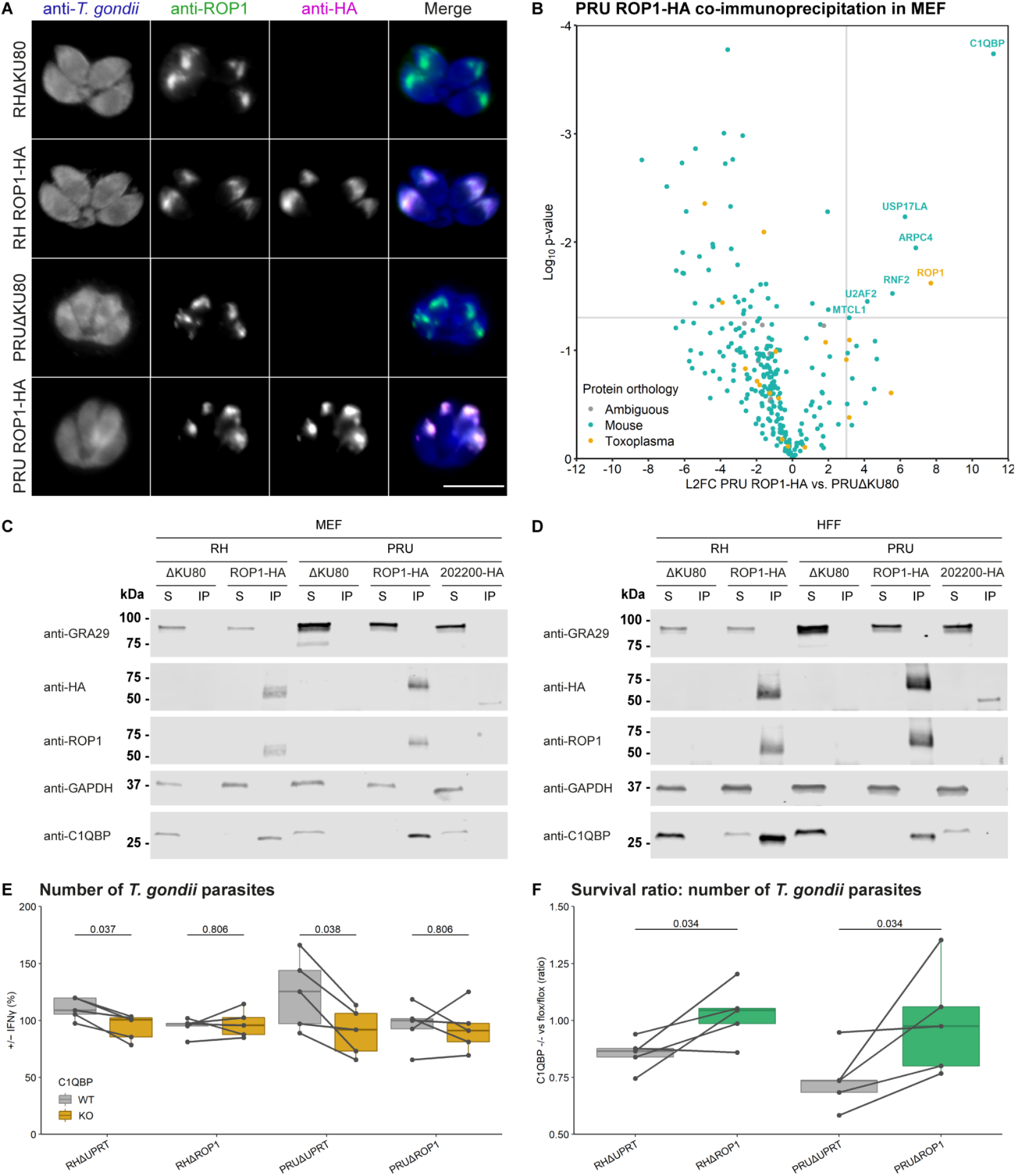
ROP1 co-immunoprecipitates with host C1QBP, which also contributes to parasite resistance to IFNγ. **A.** Immunofluorescence verification of C-terminal HA-tagging of ROP1. Scale bar = 10 μm. **B.** Enrichment of proteins that co-immunoprecipitate with ROP1. Primary MEFs were infected with PRUΔKU80 or PRU ROP1-HA parasites for 24 h, following which ROP1 was immunoprecipitated with anti-HA agarose matrix and co-immunoprecipitated proteins were identified and quantified by mass spectrometry. L2FCs were calculated from the geometric mean of the iBAQ intensities across replicates, and p-values calculated by two-sided Welch’s *t-*test. Proteins with p-value < 0.05 and L2FC > 3 are annotated. **C, D.** Co-immunoprecipitation of C1QBP with ROP1 in **C** primary MEFs and **D** HFFs infected with RH ROP1-HA and PRU ROP1-HA. S = supernatant, IP = immunoprecipitate. Note that the immunoprecipitate fraction represents 6x the relative amount of the total lysate compared to the supernatant fraction. **E.** IFNγ-dependent growth restriction of *T. gondii* in C1QBP^flox/flox^ (WT) and C1QBP^-/-^ (KO) immortalised MEFs. MEFs were stimulated with IFNγ for 24 h, infected with *T. gondii* cell lines for a further 24 h, and parasite growth quantified by automated fluorescence imaging and analysis. Parasite growth in IFNγ-stimulated BMDMs is shown as a percentage of that in unstimulated BMDMs in terms of total parasite number. p-values were calculated by paired two-sided *t-*test with Benjamini-Hochberg adjustment. **F.** Ratio of the parasite survival in C1QBP^-/-^ versus C1QBP^flox/flox^ MEFs in terms of total parasite number +/− IFNγ. p-values were calculated by paired two-sided *t-*test with Benjamini-Hochberg adjustment.

We carried out anti-HA immunoprecipitation in IFNγ-stimulated primary C57BL/6 murine embryonic fibroblasts (MEFs) infected with either PRU ROP1-HA or parental PRUΔKU80 at 24 hours post-infection. We used MEFs for this experiment due to the difficulty of obtaining large enough quantities of BMDMs and because MEFs are thought to restrict *T. gondii* through the same mechanisms as BMDMs (Saeij and Frickel 2017). Immunoprecipitated proteins were in-gel digested and identified and quantified by liquid chromatography-tandem mass spectrometry **(Supplementary Data 4)**.

As expected, ROP1 was strongly enriched in the PRU ROP1-HA-infected samples but not in the PRUΔKU80-infected samples **(Figure 4B)**. Aside from ROP1, all significantly enriched proteins were host rather than *T. gondii* proteins. Among the most highly enriched proteins were a ubiquitin-conjugating enzyme, RNF2, and a deubiquitinating enzyme, USP17LA. Therefore, we hypothesised that ROP1 may interfere with ubiquitination of the vacuole, which normally serves to recruit the guanylate-binding protein (GBP) GTPase restriction factors to the vacuole (Haldar et al. 2015). However, we did not observe any difference in the percentage of ubiquitin-decorated vacuoles of ΔROP1 and parental parasites at 3 hours post-infection in BMDMs **(Figure S6)**, suggesting that ROP1 does not impact this mechanism of restriction.

Instead, we focused on the most strongly enriched host protein in the PRU ROP1-HA samples versus PRUΔKU80: Complement Component 1q Binding Protein (C1QBP, also known as GC1QR, HABP1, p32, p33, SF2P32) **(Figure 4B)**. C1QBP is a small acidic protein which forms a homotrimer with a highly asymmetric charge distribution (Jiang et al. 1999). Intriguingly, C1QBP has been implicated as a regulator of autophagy and innate immune signalling, in addition to diverse other functions in different cellular compartments (Jiao et al. 2015; Xu et al. 2009; Waggoner et al. 2005; Petersen-Mahrt et al. 1999; Yagi et al. 2012).

To validate the interaction of ROP1 with C1QBP, we repeated the co-immunoprecipitation using both the RH and PRU strains in MEFs and HFFs and checked for enrichment of C1QBP by Western blot. We saw that in all combinations ROP1-HA pulled down C1QBP, while an unrelated HA-tagged rhoptry protein (TGME49_202200) did not **(Figure 4C & 4D)**. In the PRU strain ROP1 migrates at a higher molecular weight than in the RH strain as there is an expanded octopeptide repeat region in the middle of the protein. Since ROP1 from both strains can pull down C1QBP, we infer that this variant repeat region is likely not involved in the interaction.

### Host C1QBP facilitates parasite resistance to IFNγ

Given that ROP1 can pull down C1QBP, and that the absence of ROP1 increases susceptibility to IFNγ-mediated restriction, we hypothesised that C1QBP may have a restriction factor-like function which is inhibited by ROP1.

We observed that C1QBP localised primarily to the mitochondria **(Figure S7A)** and therefore did not see any co-localisation with ROP1. However, it has been demonstrated that there is an additional cytosolic pool of C1QBP which could putatively interact with ROP1 following injection of the rhoptry proteins or at the parasitophorous vacuole membrane (Xu et al. 2009).

To test whether the increased restriction of ΔROP1 parasites was rescued by knockout of C1QBP, we used our fluorescence microscopy growth assay to measure IFNγ-dependent restriction in immortalised MEFs derived from homozygous C1QBP^flox/flox^ C57BL/6 mice in which C1QBP had been excised by transient transfection with Cre recombinase (Yagi et al. 2012) **(Supplementary Data 5)**. Surprisingly, we observed that RHΔUPRT and PRUΔUPRT parasites were significantly more restricted in C1QBP^-/-^ MEFs than in the C1QBP^flox/flox^ MEFs, while we did not detect a significant difference for ΔROP1 parasites **(Figure 4E)**. By comparing the ratios of restriction in C1QBP^-/-^ versus C1QBP^flox/flox^ MEFs, we determined that ΔUPRT parasites were significantly more restricted than ΔROP1 parasites in C1QBP^-/-^ cells **(Figure 4F)**. We found that the major contribution to increased restriction of ΔUPRT parasites in C1QBP^-/-^ MEFs was reduced vacuole number rather than reduced vacuole size or host cell number **(Figure S8)**, as we found for ΔROP1 parasites in BMDMs and THP-1-derived macrophages. C1QBP knockout in the host cell thus phenocopies ROP1 knockout in wild-type parasites, but has no impact on ΔROP1 parasites. These data suggest that both ROP1 and C1QBP are required for full resistance to IFNγ-mediated restriction.

## DISCUSSION

We screened an expanded library of rhoptry and dense granule protein-encoding genes and identified 22 genes which contribute to *T. gondii* PRU growth *in vivo* in the mouse peritoneum, but not *in vitro*, indicating that these secreted effectors may be involved in evasion of host immune responses. Of the 235 targeted genes in this screen, we were able to determine phenotype scores for 164 genes with high confidence. This is because some protospacers had low read counts at the start of the experiment and dropped below the limit of detection over the course of the experiment. Future optimisation of CRISPR Cas9-sgRNA library preparation to achieve equal guide representation in the knockout vector pool could help to minimise drop-outs in genetic screens.

A key advantage of pooled CRISPR knockout screening is that it enables identification of virulence factors with the same function in many or all *T. gondii* strains, in contrast to genetic crosses which have been the major approach in the field until recently. By comparison to a recently published genome-wide dataset of CRISPR knockout phenotypes of the RH strain of *T. gondii* in IFNγ-stimulated BMDMs, we were able to identify a subset of effector proteins which are apparently important for immune evasion of both the PRU and RH strains, which would have been missed by genetic crosses between these strains. This comparison is likely imperfect, as different experimental models and protospacer libraries were used. Therefore, knockout screens which directly compare different strains of *T. gondii* in the same system with the same library will be an important area for future research and will provide a valuable resource to the community. Nevertheless, further study of the putative effectors important for RH and PRU infections of mice identified here, such as RASP1, GRA38, and TGME49_299780, may reveal new mechanisms of parasite virulence and subversion of the host cell.

In this work, we chose to focus on ROP1, the first-identified rhoptry protein of *T. gondii* whose function has remained mysterious for 30 years (Schwartzman 1986; Saffer et al. 1992; Ossorio, Schwartzman, and Boothroyd 1992). We demonstrated that ROP1 contributes to parasite resistance to IFNγ-induced innate immune restriction in macrophages. This phenotype likely explains the *in vivo* growth defect of ΔROP1 parasites, as macrophages are the most commonly infected cell type in acute infection in the peritoneum (Jensen et al. 2011) and IFNγ is the principle cytokine required for control of acute *T. gondii* infection *in vivo* (Suzuki et al. 1988). These results are perhaps surprising given that ROP1 knockout in the RH strain has previously been shown to have no effect on virulence in Swiss Webster mice (Soldati et al. 1995). However, the RH strain is considered “hypervirulent” in laboratory mouse strains, which can mask subtler effects on virulence which are apparent in other *T. gondii* isolates, as, for example, in the case of the Cyclase-Associated Protein (Hunt et al. 2019).

Interestingly, we found that ROP1 contributes to resistance to IFNγ-mediated restriction in both murine and human macrophages. This suggests that ROP1 counteracts an innate immune restriction mechanism which is common to both host species, although we cannot rule out a pleiotropic effect. ROP1 is protective in pre-activated immune cells, in contrast to the secreted effector IST that has been shown to protect against parasite restriction in THP-1 macrophages when interferon stimulation occurs after infection, but that is likely not protective when the host cells are pre-activated (Matta et al. 2019). To our knowledge, the dense granule chaperone GRA45 is the only secreted protein other than ROP1 known to protect against *T. gondii* clearance in human macrophages that have been pre-activated with IFNγ (Yifan Wang et al. 2020).

The rhoptries of ΔROP1 parasites have been found to have a subtly altered morphology compared to wild-type and complemented parasites, which we also observed (Soldati et al. 1995). This suggested that ROP1 may have a structural role in rhoptry function or in secretion into the host cell. However, we could not find any evidence that knockout of ROP1 affects the secretion of other rhoptry proteins, which concords with the lack of an invasion or *in vitro* growth phenotype (Soldati et al. 1995). How this ultrastructural change in the rhoptries upon deletion of ROP1 relates to the IFNγ resistance phenotype is unclear; however, the evidence suggests that IFNγ resistance is an intrinsic function of ROP1.

ROP1 from both RH and PRU parasites co-immunoprecipitates reliably with a host protein, C1QBP, from infections in both mouse and human cells. ROP1 and C1QBP are both expected to be present in small amounts in the host cell cytosol; however, we did not observe co-localisation of ROP1 with C1QBP in an infected cell and therefore cannot rule out that the interaction observed between them is an artefact of cell lysis. Both proteins are noted for highly asymmetric charge distributions which could provide the basis for any (real or artefactual) interaction: the N-terminal region of ROP1 is highly acidic and the C-terminal region is highly basic (Ossorio, Schwartzman, and Boothroyd 1992), while acidic charge is highly concentrated on one face of the C1QBP trimer (Jiang et al. 1999). The asymmetric charge distribution of ROP1 was noted by Ossorio, Schwartzman, and Boothroyd to putatively facilitate interaction with host cell components (Ossorio, Schwartzman, and Boothroyd 1992). We attempted to generate a ROP1-TurboID cell line to validate this interaction by proximity labelling; however, we found that C-terminal tagging with TurboID prevented secretion of ROP1 and localisation to the PVM, in contrast to the single HA tag that we used for co-immunoprecipitation.

How may C1QBP function in *T. gondii* restriction? Contrary to our initial hypothesis, we found that knockout of C1QBP does not rescue the phenotype of ΔROP1 parasites, as would be expected if C1QBP was a classical restriction factor. Instead, we found that C1QBP knockout made wild-type parasites more sensitive to IFNγ-mediated restriction, recapitulating the phenotype of ROP1. C1QBP has many seemingly disparate functions, so it is difficult to ascertain a possible mechanism through which its presence may benefit the parasite, although some of these described functions are potentially relevant to *T. gondii* immune evasion.

In both human and murine cells, autophagy proteins play critical roles in IFNγ-dependent parasite restriction (Saeij and Frickel 2017). C1QBP acts as a positive regulator of autophagy and mitophagy through stabilisation of ULK1 (Jiao et al. 2015). Inhibition of this function of C1QBP by binding to ROP1 would potentially account for increased parasite survival upon IFNγ stimulation, although we would not then expect C1QBP knockout MEFs to show enhanced restriction of wild-type parasites. Alternatively, activation of host autophagy through ROP1 binding to C1QBP may enhance parasite growth by providing nutrients to the parasite (Wang, Weiss, and Orlofsky 2009).

Independently of this role in autophagy, a growing body of evidence implicates C1QBP as a negative regulator of antiviral innate immunity pathways. C1QBP was found to inhibit cGAS activation in the cytosol following infection with the DNA virus HSV-1(Song et al. 2021), and to inhibit mitochondrial antiviral signaling protein (MAVS)-dependent innate immune responses upon infection with the RNA virus murine respirovirus (Sendai virus) (Xu et al. 2009). In an interesting parallel with this work, knockout/knockdown of C1QBP was found to impair viral infection in both cases. Several viral proteins have been found to interact with C1QBP, which may potentiate these inhibitory functions (Beatch et al. 2005; Waggoner et al. 2005; Lainé et al. 2003; Matthews and Russell 1998). Potentially, activation of C1QBP by binding to ROP1 could similarly inhibit these innate immune pathways.

Lacking further data we cannot determine whether the phenotypes observed for ROP1 and C1QBP are mechanistically linked. However, C1QBP knockout does not further enhance restriction of ΔROP1 parasites, in contrast to wild-type parasites, providing some evidence that the function of the two is linked and that the interaction is positive for *T. gondii* growth.

In summary, our data show that ROP1 is an important *T. gondii* effector, and that further systematic study of parasite effectors in different strains and host cell types will likely reveal yet more mechanisms of *T. gondii* immune evasion.

## METHODS

### Mice

C57BL/6 mice were bred and housed in pathogen-free conditions at the Biological Research Facility of the Francis Crick Institute in accordance with the Home Office UK Animals (Scientific Procedures) Act 1986. All work was approved by the UK Home Office (project license PDE274B7D) and the Francis Crick Institute Ethical Review Panel, and conforms to European Union directive 2010/63/EU.

### Cell culture

All cell lines were cultured at 37 °C and 5% CO_2_, and were tested monthly for *Mycoplasma spp.* contamination by PCR.

#### HFF

Primary HFFs (ATCC) were cultured in Dulbecco’s Modified Eagle’s Medium (DMEM) with 4.5 g/L glucose and GlutaMAX (Gibco) supplemented with 10% v/v heat-inactivated foetal bovine serum (FBS) (Gibco).

#### BMDM

Monocytes were isolated from the femurs of 6-12 week-old male C57BL/6 mice and differentiated into BMDMs for six days in 70% v/v RPMI 1640 medium (ATCC modification) (Gibco), 20% v/v L929 cell conditioned medium (provided by the Cell Services Science Technology Platform at the Francis Crick Institute), 10% v/v heat-inactivated FBS (Gibco), 100 U/mL penicillin-streptomycin (Gibco) and 50 μM 2-mercaptoethanol (Sigma). Following differentiation, BMDMs were cultured in the same medium without 2-mercaptoethanol.

#### THP-1

THP-1 cells were cultured in RPMI 1640 medium (Gibco) supplemented with 10% v/v FBS (Gibco). THP-1 monocytes were differentiated into macrophages with 100 ng/mL phorbol 12-myristate 13-acetate (Sigma) for 24 h, followed by a rest period without phorbol 12-myristate 13-acetate for 24 h.

#### MEF

Primary C57BL/6 MEFs (ATCC) and immortalised C57BL/6 C1QBP^flox/flox^/C1QBP^-/-^ MEFs (a gift from the lab of Dongchon Kang) were cultured in DMEM with 4.5 g/L glucose and GlutaMAX (Gibco) supplemented with 10% v/v heat-inactivated FBS (Gibco).

#### Toxoplasma gondii

All *T. gondii* tachyzoite cell lines were maintained by serial passage in HFFs. Parasites were harvested for experiments by mechanical lysis with a 27 G needle and passed through a 5 μm sterile filter. The parental lines used in this study were PRUΔHXGPRT (Donald et al. 1996), RHΔKU80 (Huynh and Carruthers 2009), and PRUΔKU80 (Fox et al. 2011). The genotypes of parasites used were verified by restriction fragment length polymorphism of the SAG3 gene (Su, Zhang, and Dubey 2006).

### CRISPR screen

#### Experimental protocol

Pooled *in vivo* CRISPR knockout screening was performed as previously described (Young et al. 2019). Briefly, ssDNA oligonucleotides encoding protospacer sequences were selected from an arrayed library using an Echo 550 Acoustic Liquid Handler (Labcyte). Five protospacer sequences were selected per target gene, and dispensed in triplicate. The ssDNA oligonucleotides were integrated into a pCas9-GFP-T2A-HXGPRT::sgRNA vector (Young et al. 2019) by pooled Gibson assembly.

The resulting plasmid pool was linearised and transfected into 10^7^ PRUΔHXGPRT tachyzoites using the Amaxa 4D Nucleofector system (Lonza) with buffer P3 and pulse code EO-115. After 24 h recovery, transfected parasites were selected in HFFs for integration of the plasmid into the genome with 25 μg/mL mycophenolic acid (Sigma) and 50 μg/mL xanthine (Sigma). Following selection, five mice were infected by intraperitoneal injection with 200,000 parasites each, as determined by plaque assay. After five days, parasites were recovered by peritoneal lavage and cultured in HFFs for one passage.

Genomic DNA was extracted from a sample of the parasite population following *in vitro* drug selection, from the leftover mouse inoculum, and from the five *ex vivo* populations using the DNeasy Blood and Tissue Kit (Qiagen). Illumina sequencing libraries were prepared by nested PCR amplification of the protospacer sequences from the parasite genomic DNA and the plasmid pool using primers 1-11. The libraries were sequenced on a HiSeq 4000 platform (Illumina) with 100 bp paired-end reads to a minimum depth of 7.5 million reads per sample (approximately 5000x coverage of the protospacer pool).

#### Data analysis

Following demultiplexing, the reads were trimmed and aligned to a reference of protospacer sequences using a custom perl script. Subsequent analysis was carried out using R v4.0.1 (https://www.r-project.org/) with packages tidyverse v1.3.1, qvalue v2.22.0, ggrepel v0.9.1 and scales v1.1.1.

Protospacers with fewer than 50 raw reads in every sample were removed from the analysis and remaining counts normalised using the median of ratios method (Anders and Huber 2010). Genes with fewer than three protospacers remaining were then removed from the analysis.

For each gene, the median *in vitro* L2FC was calculated from the normalised counts of the protospacers targeting that gene in the drug-selected parasite population and the plasmid pool. The median *in vivo* L2FC was calculated using the geometric mean of the normalised counts in the *ex vivo* parasite populations and the normalised counts in the inoculum. Genes in the top 5^th^ percentile of median absolute deviation of the *in vitro* or *in vivo* L2FCs were removed from the analysis.

In addition, for each gene an *in vitro* and *in vivo* p-value calculated by paired two-sided t-test on the log_2_-transformed normalised counts and adjusted to correct for local false discovery rate (FDR) using the qvalue R package. The median L2FCs and FDR-adjusted q-values were used to calculate a DISCO score for each gene as:

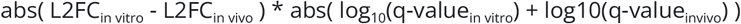

### Generation of *T. gondii* cell lines

#### Knockouts

Inverse PCR was used to introduce a protospacer targeting the CDS of either ROP1 or UPRT to a pCas9-GFP::sgRNA plasmid using primers 12-14. For ROP1, A Pro^GRA1^-mCherry-T2A-HXGPRT-Ter^GRA2^ construct was amplified from a template plasmid (Young et al. 2019) using primers 15 and 16 to induce 40 bp homology arms to the 5’ and 3’ UTRs of ROP1. For UPRT, the above construct was amplified using primers 17 and 18 and integrated into BamHI/PacI-digested (NEB) pUPRT plasmid by Gibson assembly. 15 μg of homology repair template (purified PCR product or linearised pUPRT-mCherry-HXGPRT) was co-transfected with 15 μg pCas9 plasmid targeting the gene of interest into the RHΔKU80 and PRUΔKU80 lines using the Amaxa 4D Nucleofector system (Lonza) as above. After 24 h recovery, transfected parasites were selected with 25 μg/mL mycophenolic acid and 50 μg/mL xanthine for at least six days before single-cell cloning by serial dilution. Integration of the mCherry-HXGPRT cassette repair template was verified by PCR with primers 19-22.

#### Complementation

The ROP1 CDS together with 1000 bp upstream of the start codon was amplified from RHΔKU80 and PRUΔKU80 genomic DNA using primers 23-25. The backbone of the pUPRT plasmid was amplified with primers 26 and 27 and assembled with the ROP1 inserts by Gibson assembly. 15 μg of pUPRT-RH/PRU-ROP1-HA plasmid was linearised and transfected together with 15 μg pCas9 plasmid targeting UPRT into the RHΔROP1 and PRUΔROP1 lines. After 24 h recovery, transfected parasites were selected with 5 μM 5′-fluo-2′-deoxyuridine for at least six days before single-cell cloning by serial dilution. Integration of the pUPRT-RH/PRU-ROP1-HA plasmids into the UPRT locus was verified by PCR using primers 21 and 28.

#### HA tagging

Inverse PCR was used to introduce a protospacer targeting the 3’ UTR of ROP1 into the pCas9-GFP::sgRNA plasmid using primers 12 and 29. Primers 30 and 31 were used to amplify an in-frame HA-Ter^GRA2^::Pro^DHFR^-HXGPRT-Ter^DHFR^ construct from a template plasmid, introducing 40 bp homology arms to the 3’ end of the ROP1 CDS. 15 μg each of pCas9 plasmid and purified PCR product were co-transfected as above into the RHΔKU80 and PRUΔKU80 lines. Selection with mycophenolic acid and xanthine and cloning were carried out as above. Integration of the HA-tag repair construct was verified with primers 32 and 33.

### ROP1 immunofluorescence assays

HFFs were grown to confluence in an 8-well μ-slide (Ibidi) and infected with *T. gondii* strains for 24 h. The slides were fixed with 4% w/v formaldehyde (Sigma) in phosphate-buffered saline (PBS) (Sigma). The cells were permeabilised with 0.2% v/v Triton X-100 (Sigma) for 15 min or 1 min and blocked with 2% w/v bovine serum albumin (Sigma) for 1 h. The cells were stained with 1:500 rat anti-HA (Roche #11867423001), followed by 1:1000 goat anti-rat 594 (Invitrogen #A11007), followed by a mixture of 1:500 mouse anti-ROP1 (Abnova #MAB17504) and 1:1000 rabbit anti-T. gondii (Abcam #ab138698), and finally with a mixture of 1:1000 goat anti-mouse 488 (Invitrogen #A11029), 1:1000 goat anti-rabbit 647 (Invitrogen #A21244), and 5 μg/mL DAPI (Sigma), each for 1h at room temperature. Images were acquired on a Nikon Ti-E inverted widefield fluorescence microscope with a Nikon CFI APO TIRF 100x/1.49 objective and Hamamatsu C11440 ORCA Flash 4.0 camera running NIS Elements (Nikon).

### ROP1 Western blotting

Parasites were purified from host cell material by syringe-lysis, filtering and washing in PBS, then lysed in RIPA buffer (Pierce) supplemented with 2x cOmplete Mini EDTA-free Protease Inhibitor Cocktail (Roche). 10 μg protein per sample was boiled for 5 min in sample loading buffer and separated by SDS-PAGE using the Mini-PROTEAN electrophoresis system (Bio-Rad). Proteins were transferred to a nitrocellulose membrane using the Trans-Blot Turbo transfer system (Bio-Rad), blocked in 2% w/v skim milk powder, 0.1% v/v Tween 20 in PBS for 1 h at room temperature, then incubated with primary antibodies in blocking buffer overnight at 4 °C. Primary antibodies used were 1:1000 mouse anti-ROP1 (Abnova #MAB17504), 1:1000 rat anti-HA (Roche #11867423001), and 1:200 mouse anti-*T. gondii* (Santa Cruz #SC-52255). Blots were stained with secondary antibodies for 1 h at room temperature: 1:10,000 goat anti-rat IRDye 680LT (Li-Cor #925-68029) and 1:10,000 goat anti-mouse IRDye 800CW (Li-Cor #925-32210). Blots were visualised using an Odyssey CLx scanner (Li-Cor).

### Plaque assays

100 parasites were inoculated onto a T25 flask of confluent HFFs and left undisturbed for seven days, following which the cells were stained with 0.5% w/v crystal violet (Sigma), 0.9% w/v ammonium oxalate (Sigma), 20% v/v methanol in distilled water.

### IFNγ restriction assays

For BMDMs, 75,000 cells per well were seeded in a 96-well μ-plate (Ibidi). For THP-1-derived macrophages, 75,000 THP-1 monocytes were seeded per well and differentiated into macrophages as above. For MEFs, 10,000 cells were seeded per well and grown to confluence prior to infection. BMDMs and MEFs were stimulated with 10 ng/mL (~100 U/mL) recombinant mouse IFNγ (Gibco) for 24 h prior to infection or left unstimulated. THP-1-derived macrophages were stimulated with 50 ng/mL (~100 U/mL) recombinant human IFNγ (BioTechne) for 24 h prior to infection or left unstimulated. The plates were infected with parasite lines at an MOI of 0.3 for 24 h, with at least three wells for each line with and without IFNγ. The plates were fixed with 4% w/v formaldehyde for 15 min and stained with 5 μg/mL DAPI and 5 μg/mL CellMask deep red plasma membrane stain (Invitrogen) for 1 h at room temperature. Biological replicates were carried out on different days with independently prepared host cells.

The plates were imaged on an Opera Phenix High-Content Screening System (PerkinElmer) with a 40x/1.1 NA water immersion objective. 25 fields of view with 3-5 focal planes (depending on the host cell type) were imaged per well. Analysis was performed in Harmony v5 (PerkinElmer) on a maximum projection of the planes. Image acquisition parameters and analysis sequence are detailed in **Supplementary Data 7**. For each well, the total number of host cell nuclei and *T. gondii* vacuoles in the captured fields of view was determined by thresholding on the DAPI and mCherry signal. The number of parasite nuclei in each vacuole was determined based on DAPI signal to define the total number of parasites the the captured fields of view and the mean number of parasites per vacuole in each well. For each *T. gondii* line, IFNγ-mediated restriction was calculated as the median tachyzoite number/vacuole number/vacuole size/host cell number in the IFNγ-stimulated wells as a percentage of the median in the unstimulated wells. Differences between strains were tested by paired two-sided *t*-test with Benjamini-Hochberg adjustment.

### Transmission electron microscopy

Confluent HFFs grown on glass coverslips were infected with *T. gondii* lines for 24 h, fixed with 2.5% glutaraldehyde, 4% formaldehyde in 0.1 M phosphate buffer (PB) for 30 min and transferred to a BioWave Pro+ microwave for processing (Pelco; Agar Scientific). The microwave program used is detailed in **Supplementary Data 8**. The cells were washed with PB twice on the bench and twice in the microwave 250 W for 40 s, stained with 1% reduced osmium for 14 min under vacuum (with/without 100 W power at 2 min intervals), and then washed twice on the bench and twice in the microwave with PB. A further stain with 1% tannic acid for 14 min (with/without 100 W power at 2 min intervals under vacuum) was followed by a quench with 1% sodium sulfate at 250 W for 2 min under vacuum and bench and microwave washes in water (as for PB). The blocks were then dehydrated in a graded ethanol series of 70%, 90%, and 100%, each performed twice at 250 W for 40 s. Exchange into Epon resin (Taab Embed 812) was performed with 50% resin in ethanol, followed by three 100% resin steps, each at 250 W for 3 min, with 30 s vacuum cycling. Finally, the samples were baked for 24 h at 60°C. 80 nm sections were stained with lead citrate and imaged in a JEM-1400 FLASH transmission electron microscope (JEOL).

### Rhoptry secretion assays

#### ROP16-mediated phosphorylation of STAT6

T25 flasks of confluent HFFs were infected with 1 million parasites for 2 h, after which the HFFs were dissociated and fixed with methanol for 10 minutes. The cells were stained with 1:200 rabbit anti-phospho-STAT6 (Cell Signaling #56554) and 1:200 mouse anti-*T. gondii* (Santa Cruz #SC-52255) overnight at 4 °C, followed by 1:1000 goat anti-rabbit 488 (Invitrogen #A11008), 1:1000 goat anti-mouse 594 (Invitrogen #A11005), and 5 μg/mL DAPI for 1 h at room temperature. Data were collected on an LSR II flow cytometer (BD) running FACSDiva v9 (BD) and analysed with FlowJo v10 (www.flowjo.com). The median anti-phospho-STAT6 signal in the infected cells was determined for each sample, and the median technical replicate taken to represent the biological replicate. The data were scaled to RHΔKU80 = 1 AU and differences between strains tested by two-sided Wilcoxon rank sum test with Bonferroni correction.

#### ROP17-dependent induction of cMyc

HFFs were grown to confluence in an 8-well μ-slide (Ibidi) and serum starved for 24 h before infection in 0.1% FBS medium. Each well was infected with 40,000 parasites for 24 h in 0.1% FBS medium before fixation with 4% w/v formaldehyde for 15 min, permeabilisation with 0.2% v/v Triton X-100 for 15 min, and blocking with 2% w/v BSA for 1 h. The cells were stained with 1:800 rabbit anti-cMyc (Cell Signaling #5605) and 1:200 mouse anti-*T. gondii* (Santa Cruz #SC-52255) for 2 h at room temperature, followed by 1:1000 goat anti-rabbit 488 (Invitrogen #A11008), 1:1000 goat anti-mouse 594 (Invitrogen #A11005), and 5 μg/mL DAPI for 1 h at room temperature. Images were acquired on a Nikon Ti-E inverted widefield fluorescence microscope with a Nikon Plan APO 40x/0.95 objective and Hamamatsu C11440 ORCA Flash 4.0 camera running NIS Elements (Nikon) and analysed using ImageJ (Schneider, Rasband, and Eliceiri 2012). The median cMyc fluorescence intensity in each nucleus was determined and the median nucleus taken as representative of a replicate. The median background cMyc fluorescence intensity was subtracted, and the data normalised to RHΔKU80 = 1 AU for each biological replicate. Differences between strains were tested by two-sided Wilcoxon rank sum test with Bonferroni correction.

#### ROP18-dependent inhibition of IRGB6 recruitment

75,000 BMDMs per well were seeded in a 96-well μ-plate (Ibidi) and stimulated with 10 ng/mL (~100 U/mL) recombinant mouse IFNγ (Gibco) for 24 h prior to infection. The BMDMs were infected with parasite strains with an MOI of 0.3 for 1 h, fixed with 4% w/v formaldehyde for 15 min, permeabilised with 0.1% w/v saponin (Sigma) for 15 minutes and blocked with 2% w/v BSA for 1 h. The plate was stained with 1:4000 rabbit anti-IRGB6 (a gift from the lab of Jonathan Howard) for 1 h at room temperature, followed by 1:1000 goat anti-rabbit 488 (Invitrogen #A11008), 5 μg/mL DAPI, and 5 μg/mL CellMask Deep Red plasma membrane stain for 1 h at room temperature. Images were acquired on an Opera Phenix High-Content Screening System (PerkinElmer) as above, and analysed in Harmony v5. Image acquisition parameters and analysis sequence are detailed in **Supplementary Data 7**. Vacuoles were counted as recruited if the median anti-IRGB6 intensity in a 6 pixel-wide ring around the vacuole (defined by parasite-expressed mCherry signal) was more than 2.3-2.6x (depending on the maximum signal intensity in the replicate) higher than the median anti-IRGB6 signal in the rest of the infected cell. For each well the % IRGB6-recruited vacuoles was determined, and the median % recruitment per strain taken as representative of a biological replicate. Differences between strains were determined by paired two-sided *t-*test with Bonferroni correction.

### Co-immunoprecipitation

Primary MEFs/HFFs were grown to confluence in T175 flasks and infected with 5 million parasites per flask for 24 h. The flasks were washed twice with chilled PBS and lysed in 1 mL IP buffer on ice (50 mM Tris, 150 mM NaCl, 0.2% v/v Triton X-100, 2x cOmplete Mini EDTA-free Protease Inhibitor Cocktail, pH 7.5).

#### Co-immunoprecipitation-mass spectrometry

For mass spectrometry analysis, the lysates were incubated with 40 uL per sample of Pierce anti-HA agarose matrix (Thermo) overnight at 4 °C, following which the matrix was washed three times with IP buffer and the bound proteins eluted with 30 μL 3x Sample Loading Buffer (NEB) at 95 °C for 10 min. Samples were prepared for LC-MS/MS analysis by in-gel tryptic digestion. Briefly, the eluted proteins were run 1 cm into a NuPAGE 10% Bis-Tris gel (Invitrogen) and stained with Coomassie Brilliant Blue. The gel was cut into 1 mm cubes, destained using 50% ethanol, 50 mM ammonium bicarbonate, and dehydrated with 100% ethanol. Proteins were then simultaneously reduced and alkylated with 10 mM tris(2-carboxyethyl)phosphine and 40 mM chloroacteamide in water at 70 °C for 5 min. The gel cubes were washed in 50% ethanol, 50 mM ammonium bicarbonate and dehydrated as above. Proteins were digested with 250 ng of mass spectrometry-grade trypsin (Thermo) in 50 mM HEPES, pH 8, at 37 °C overnight. Peptides were extracted from the gel into acetonitrile and dried by vacuum centrifugation.

Digested samples were solubilised in 0.1% formic acid and loaded onto Evotips (Evosep), according to the manufacturer’s instructions. Following a wash with aqueous acidic buffer (0.1% formic acid in water), samples were loaded onto an Evosep One system coupled to an Orbitrap Fusion Lumos (ThermoFisher Scientific). The Evosep One was fitted with a 15 cm column (PepSep) and a predefined gradient for a 44-minute method was employed. The Orbitrap Lumos was operated in data-dependent mode (1 second cycle time), acquiring IT HCD MS/MS scans in rapid mode after an OT MS1 survey scan (R=60,000). The MS1 target was 4E5 ions whereas the MS2 target was 1E4 ions. The maximum ion injection time utilized for MS2 scans was 300 ms, the HCD normalized collision energy was set at 32 and the dynamic exclusion was set at 15 seconds.

Acquired raw files were processed with MaxQuant v1.5.2.8 (Cox and Mann 2008). Peptides were identified from the MS/MS spectra searched against *Toxoplasma gondii* (ToxoDB) and *Mus musculus* (UniProt) proteomes using the Andromeda search engine (Cox et al. 2011). Methionine oxidation, acetylation (N-term), and deamidation (NQ) were selected as variable modifications whereas cysteine carbamidomethylation was selected as a fixed modification. The enzyme specificity was set to trypsin with a maximum of two missed cleavages. The precursor mass tolerance was set to 20 ppm for the first search (used for mass re-calibration) and to 4.5 ppm for the main search. The datasets were filtered on posterior error probability (PEP) to achieve a 1% false discovery rate on protein, peptide and site level. Other parameters were used as pre-set in the software. “Unique and razor peptides” mode was selected to allow identification and quantification of proteins in groups (razor peptides are uniquely assigned to protein groups and not to individual proteins). Intensity-based absolute quantification (iBAQ) in MaxQuant was performed using a built-in quantification algorithm (Cox and Mann 2008) enabling the “Match between runs” option (time window 0.7 minutes) within replicates.

MaxQuant output files were processed with Perseus, v1.5.0.9 (Tyanova et al. 2016) and Microsoft Office Excel 2016 **(Supplementary Data 4)**. Data were filtered to remove contaminants, protein IDs originating from reverse decoy sequences and only identified by site. iBAQ intensities and the total intensity were log_2_ and log_10_ transformed, respectively. Samples were grouped according to sample type (PRUΔKU80 or ROP1-HA) and the iBAQ intensities were filtered for the presence of two valid values in at least one group. Next, missing values were imputed from the normal distribution in order to generate log_2_ fold-changes (L2FCs) between tested conditions and perform statistical analysis (Welch’s *t*-test, p < 0.05, −3 > L2FC > 3). The L2FC threshold was set at three times the median absolute deviation.

The mass spectrometry proteomics data have been deposited to the ProteomeXchange Consortium via the PRIDE (Perez-Riverol et al. 2019) partner repository with the dataset identifier PXD032319.

#### Co-immunoprecipitation-Western blot

For Western blotting analysis, the lysate was incubated with 30 μL Pierce anti-HA magnetic beads (Thermo) overnight at 4 °C, following which the beads were washed three times with IP buffer and the bound proteins eluted with 30 μL 3x Sample Loading Buffer (NEB) at 95 °C for 10 min. Eluted proteins were separated by SDS-PAGE and transferred to a nitrocellulose membrane as above. The membrane was blocked with 2% w/v skim milk powder, 0.1% v/v Tween 20 in PBS for 1 h at room temperature, then incubated with primary antibodies in blocking buffer overnight at 4 °C. Primary antibodies used were: 1:1000 rat anti-HA (Roche #11867423001), 1:1000 mouse anti-ROP1 (Abnova #MAB17504), 1:1000 rabbit anti-C1QBP (Abcam #ab270032), 1:10,000 rabbit anti-GAPDH (Proteintech #10494-1-AP), and 1:1000 rabbit anti-GRA29 (Young et al. 2019). Blots were stained with secondary antibodies for 1 h at room temperature: 1:10,000 goat anti-mouse IRDye 680LT (Li-Cor #925-68020), 1:10,000 goat anti-rat IRDye 800CW (Li-Cor #925-32219), 1:10,000 donkey anti-rabbit IRDye 680LT (Li-Cor #925-68023), and 1:10,000 donkey anti-rabbit IRDye 800CW (Li-Cor #925-32213). Blots were visualised using an Odyssey CLx scanner (Li-Cor).

### Vacuole ubiquitination assay

150,000 BMDMs per well were seeded in an 8-well μ-slide and stimulated with 10 ng/mL (~100 U/mL) recombinant mouse IFNγ for 24 h prior to infection. The BMDMs were infected with parasite strains at an MOI of 0.3 for 3 h, then washed and fixed with 4% w/v formaldehyde for 15 min. Prior to permeabilisation, the cells were blocked with 2% w/v BSA for 1 h and extracellular parasites were stained with 1:1000 rabbit anti-*T. gondii* (Abcam #ab138698) for 1 h at room temperature followed by 1:1000 goat anti-rabbit 405 (Invitrogen #A31556) for 1 h at room temperature. The cells were then permeabilsed with 0.2% v/v Triton X-100 for 15 minutes, blocked again with 2% w/v BSA for 1 h, stained with 1:200 mouse anti-ubiquitinylated proteins (Sigma #04-263) overnight at 4 °C, and finally stained with 1:1000 goat anti-mouse 488 (Invitrogen #A11029) for 1 h at room temperature. Nine tiled fields of view were captured for each well on a Nikon Ti-E inverted widefield fluorescence microscope as above. The images were blinded, and the percentage of ubiquitinated vacuoles was determined manually using ImageJ, excluding *T. gondii* cells which were positive for extracellular staining. A median of 290 vacuoles were analysed per strain per replicate. Differences between strains were determined by two-sided *t*-test with Bonferroni correction.

## ACKNOWLEDGEMENTS

We thank all members of the Treeck laboratory for critical discussions. We thank Rachael Instrell and Becky Saunders (High-Throughput Screening Science Technology Platform, The Francis Crick Institute, London, United Kingdom) for assistance with sgRNA library preparation. We thank Takeshi Uchiumi and Dongchon Kang (Kyushu University, Fukuoka, Japan) for providing the C1QBP^flox/flox^ and C1QBP^-/-^ MEFs. We thank members of the Advanced Sequencing Facility, Biological Research Facility, and Cell Services Science Technology Platforms at the Francis Crick Institute for support.

This work was supported by funding to MT from the Francis Crick Institute which receives its core funding from Cancer Research UK (FC001189), the UK Medical Research Council (FC001189), and the Wellcome Trust (FC001189). The Science Technology Platforms at the Francis Crick Institute receive funding from Cancer Research UK (FC001999), The UK Medical Research Council (FC001999) and the Wellcome Trust (FC001999). For the purpose of Open Access, the authors have applied a CC BY public copyright licence to any Author Accepted Manuscript version arising from this submission.

## AUTHOR CONTRIBUTIONS

Conceptualisation: SB, JW, JY, MT. Investigation: SB, FT, EL, JW, OS, MB, MR, JY. Formal analysis: SB, MB. Visualisation: SB. Supervision: JY, MT. Project administration: MT. Funding acquisition: MT. Writing - original draft: SB, MT. Writing - review and editing: all authors

## SUPPLEMENTARY FIGURES

**Figure S1:**
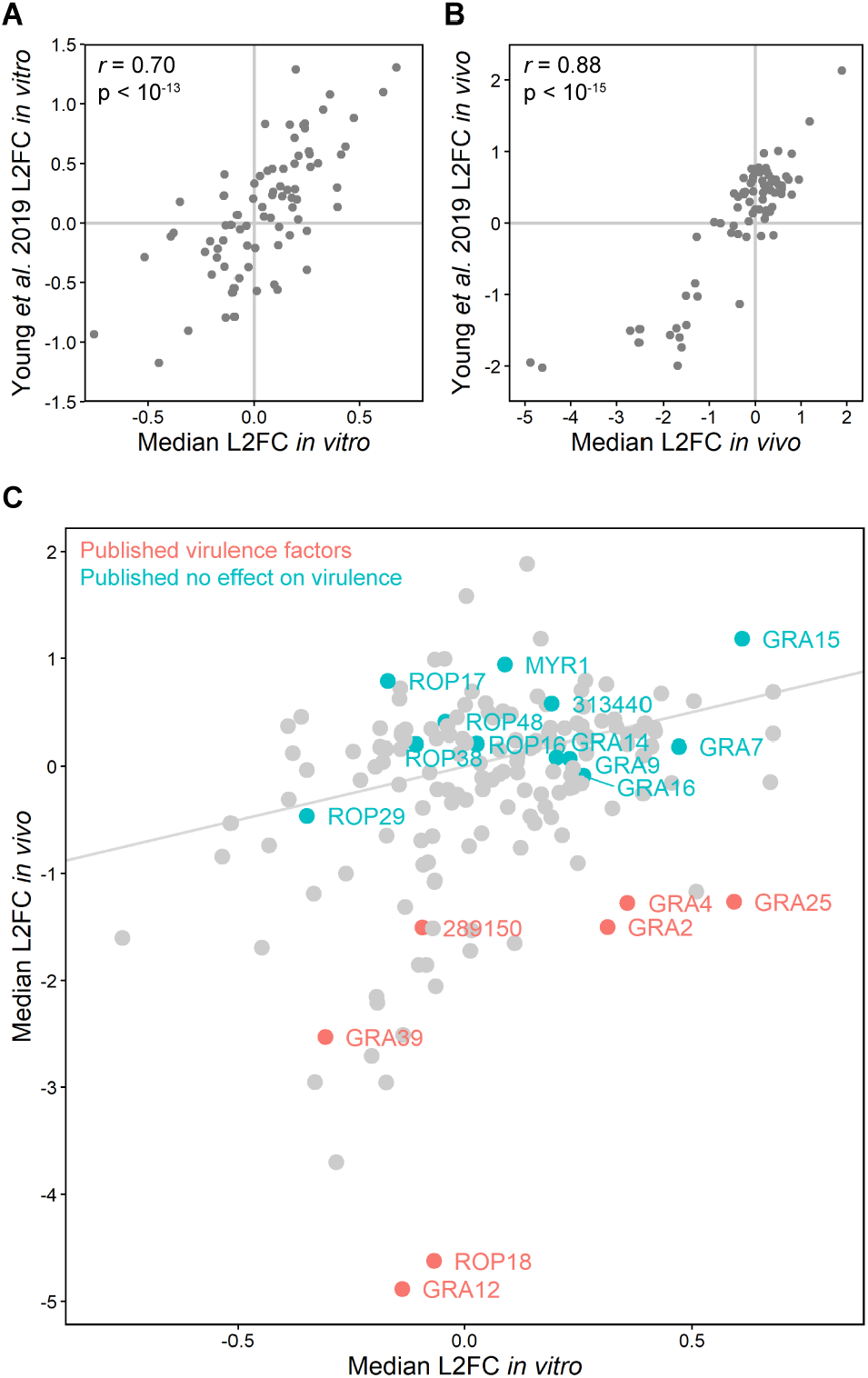
CRISPR screen phenotypes are reproducible and accurately separate known virulence factors. **A, B.** Correlation between **A** *in vitro* L2FCs and **B** *in vivo* L2FCs from this study and from (Young et al. 2019). *r* = Pearson’s product-moment correlation coefficient. **C.** Scatter plot of median L2FCs for each gene *in vitro* and *in vivo*. Genes which have previously been tested for an effect on virulence in the PRU strain of *T. gondii* are labelled.

**Figure S2:**
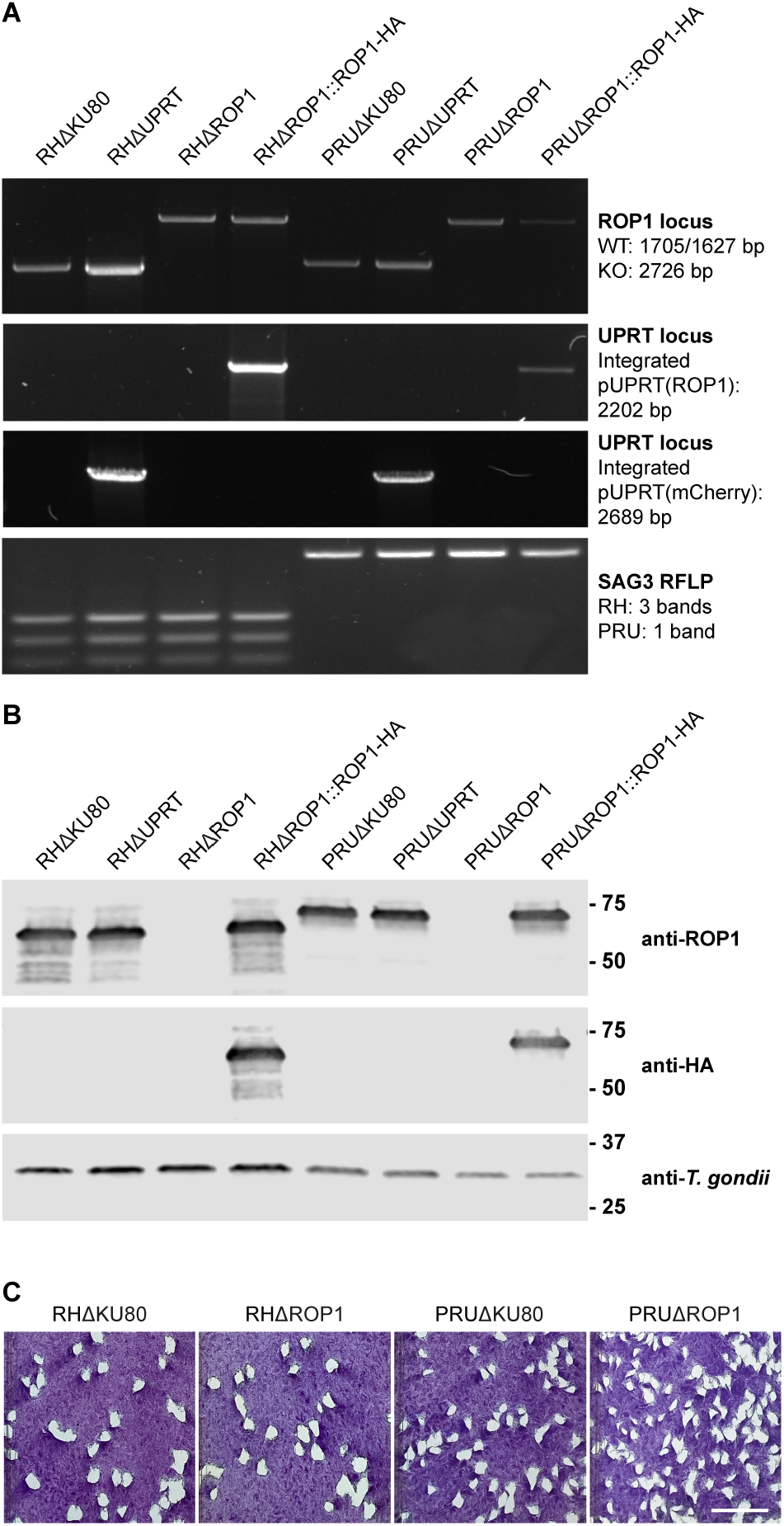
Verification of *T. gondii* knockout cell lines by PCR and Western blot. **A.** Verification of correct integration of knockout and complementation constructs by diagnostic PCR and verification of strain genotype by restriction fragment length polymorphism (RFLP) of the SAG3 gene (Su, Zhang, and Dubey 2006). Knockouts were obtained by integration of an mCherry-T2A-HXGPRT linear PCR cassette facilitated by co-transfection with a Cas9-sgRNA plasmid targeting the gene of interest. For ROP1 complementation, the ROP1 coding sequence and native promoter were cloned from RHΔKU80 or PRUΔKU80 genomic DNA into the pUPRT vector, adding a single C-terminal HA tag. Linearised pUPRT(ROP1-HA) plasmids were integrated by double homologous recombination following co-transfection with a Cas9-sgRNA plasmid targeting the UPRT locus. Clonal *T. gondii* cell lines were obtained by limiting dilution. **B.** Verification of ROP1 and ROP1-HA expression by Western blot. **C.** Plaques formed by RHΔKU80, RHΔROP1, PRUΔKU80 and PRUΔROP1 parasites after seven days’ growth in a monolayer of HFFs. Scale bar = 1 cm.

**Figure S3:**
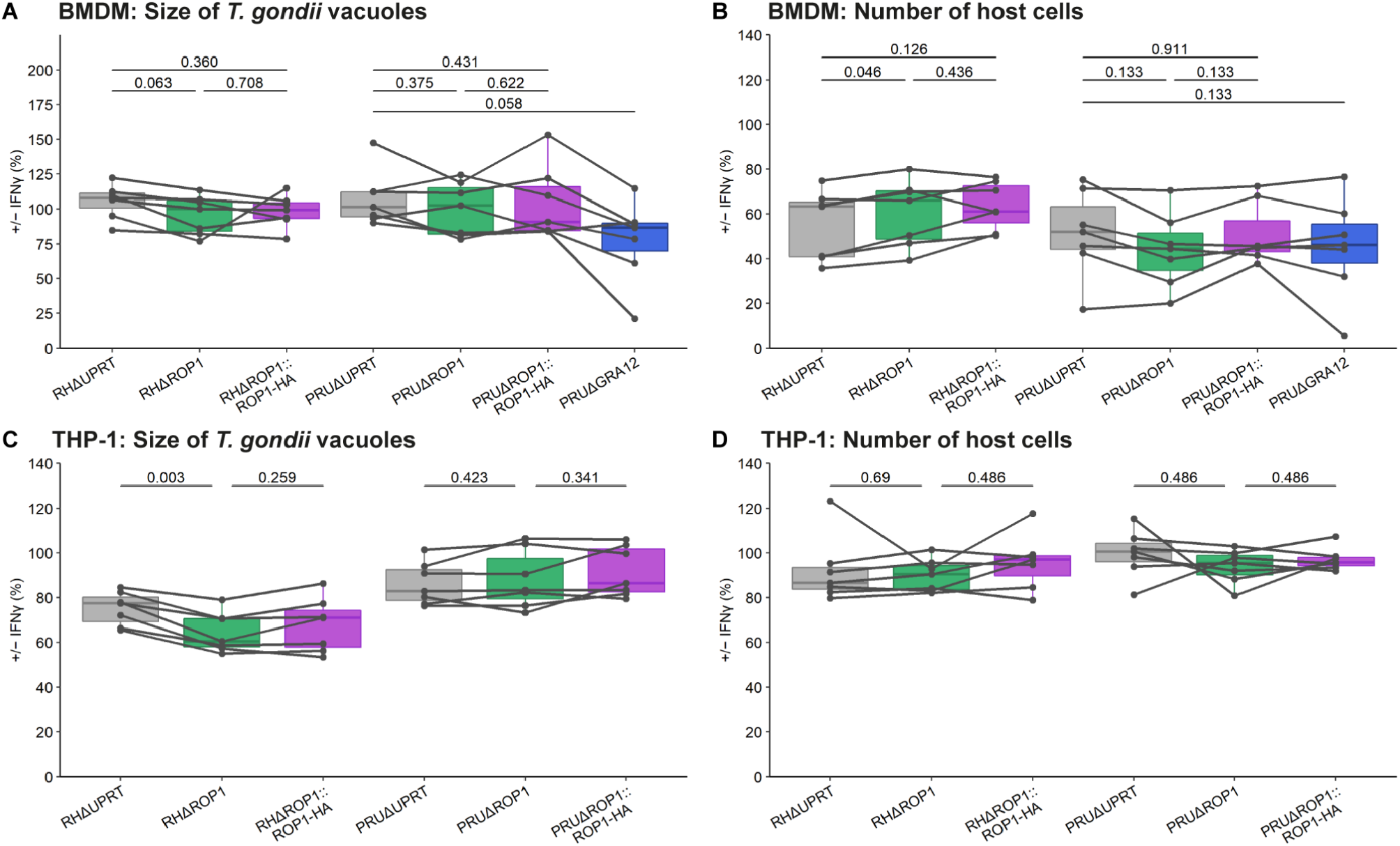
ROP1 does not affect vacuole size or host cell death. **A, B.** IFNγ-dependent growth restriction of *T. gondii* in BMDMs. BMDMs were stimulated with IFNγ for 24 h, infected with *T. gondii* cell lines for a further 24 h and parasite growth quantified by automated fluorescence imaging and analysis. **A** *T. gondii* vacuole size (mean parasites per vacuole) in IFNγ-stimulated BMDMs is shown as a percentage of the size in unstimulated BMDMs. **B** The number of IFNγ-stimulated host BMDM nuclei is shown as a percentage of the number of unstimulated BMDM nuclei. p-values were calculated by paired two-sided *t-*test with Benjamini-Hochberg adjustment. **C, D.** IFNγ-dependent growth restriction of *T. gondii* in THP-1-derived macrophages. Differentiated THP-1 macrophages were stimulated with IFNγ, infected, and parasite growth quantified as above. **C** *T. gondii* vacuole size (mean parasites per vacuole) in IFNγ-stimulated THP-1 macrophages is shown as a percentage of the size in unstimulated macrophages. **D** The number of IFNγ-stimulated host THP-1 macrophage nuclei is shown as a percentage of the number of unstimulated macrophage nuclei. p-values were calculated by paired two-sided *t-*test with Benjamini-Hochberg adjustment.

**Figure S4:**
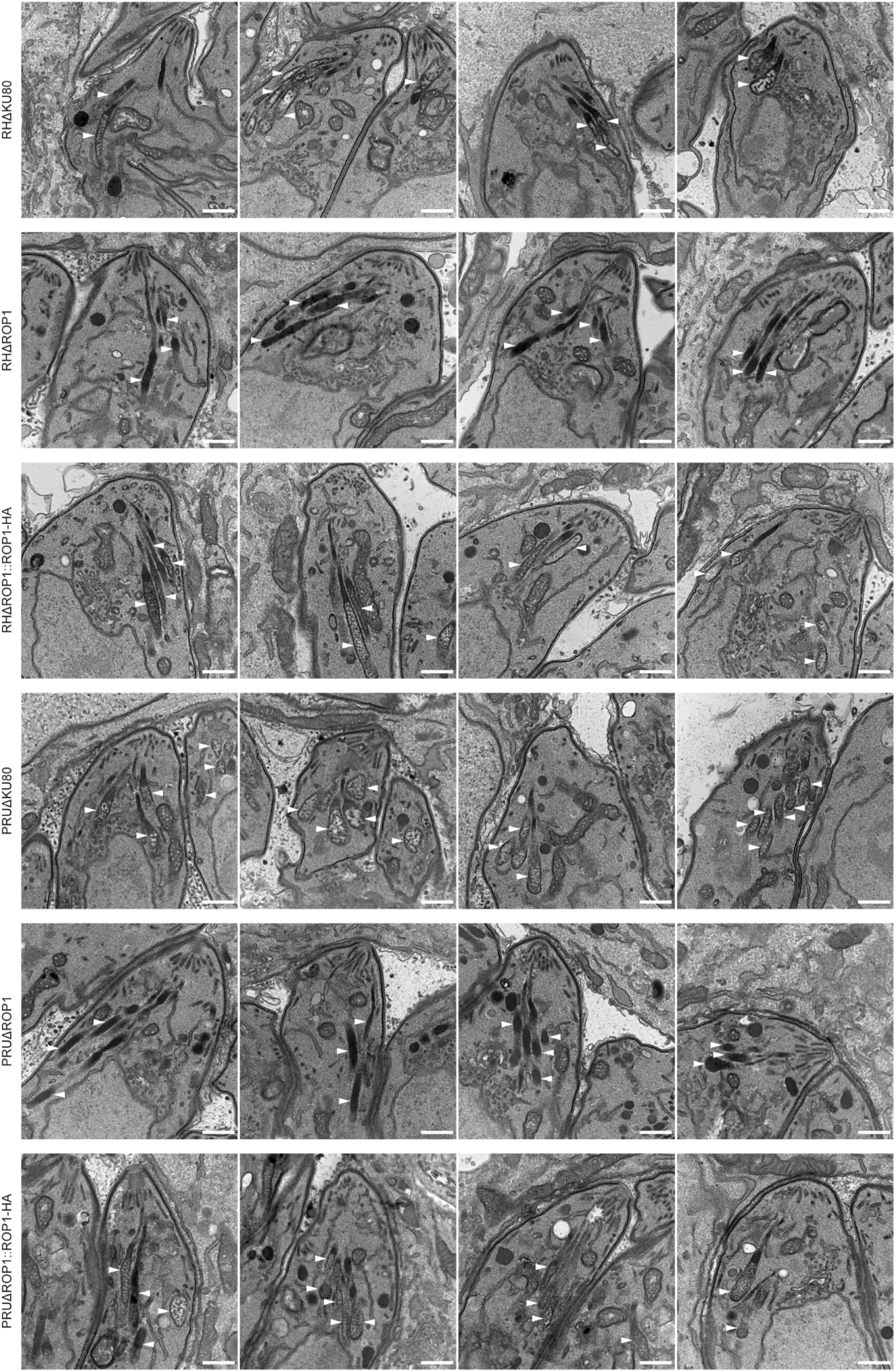
Additional rhoptry TEM images. White arrowheads indicate rhoptries. Scale bar = 500 μm.

**Figure S5:**
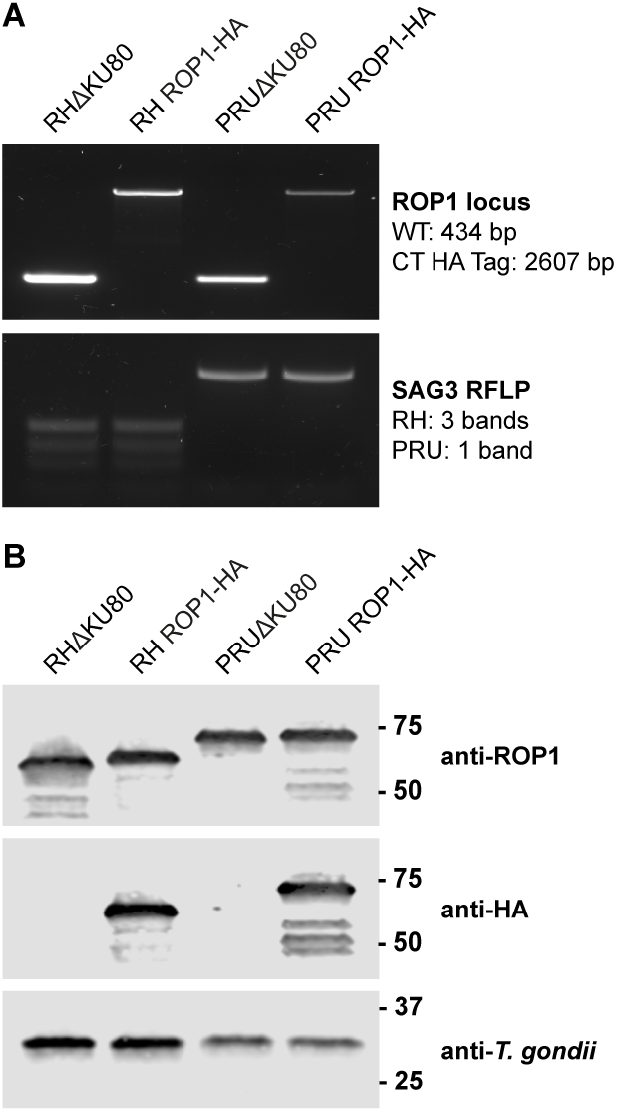
Verification of *T. gondii* C-terminal tagged cell lines by PCR and Western blot. **A.** Verification of correct integration of C-terminal HA-tagging construct by diagnostic PCR and verification of strain genotype by restriction fragment length polymorphism (RFLP) of the SAG3 gene (Su, Zhang, and Dubey 2006). HA-tagged cell lines were obtained by double homologous recombination with an HA-HXGPRT linear PCR cassette facilitated by co-transfection with a Cas9-sgRNA plasmid targeting the 3’ UTR of ROP1. Clonal *T. gondii* cell lines were obtained by limiting dilution. **B.** Verification of ROP1 and ROP1-HA expression by Western blot.

**Figure S6:**
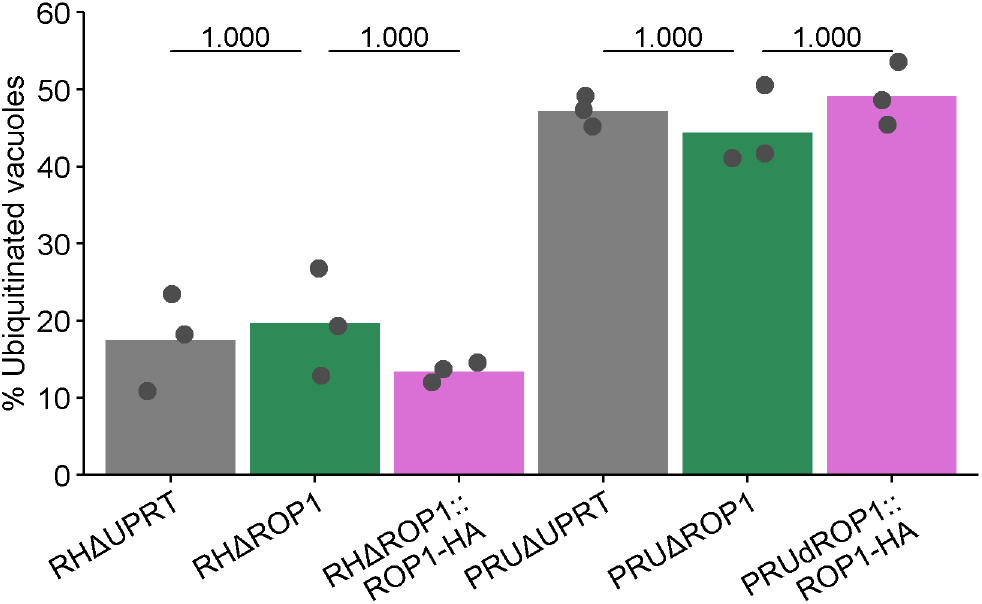
ROP1 does not affect vacuole ubiquitination. Percentage of ubiquitinated vacuoles. BMDMs were stimulated with 100 U/mL IFNγ for 24 h, infected for 3 h, fixed, and stained with an anti-ubiquitinylated proteins antibody. The percentage of ubiquitinated vacuoles was quantified manually from blinded immunofluorescence microscopy images. p-values were calculated by paired two-sided *t-*test with Bonferroni correction.

**Figure S7:**
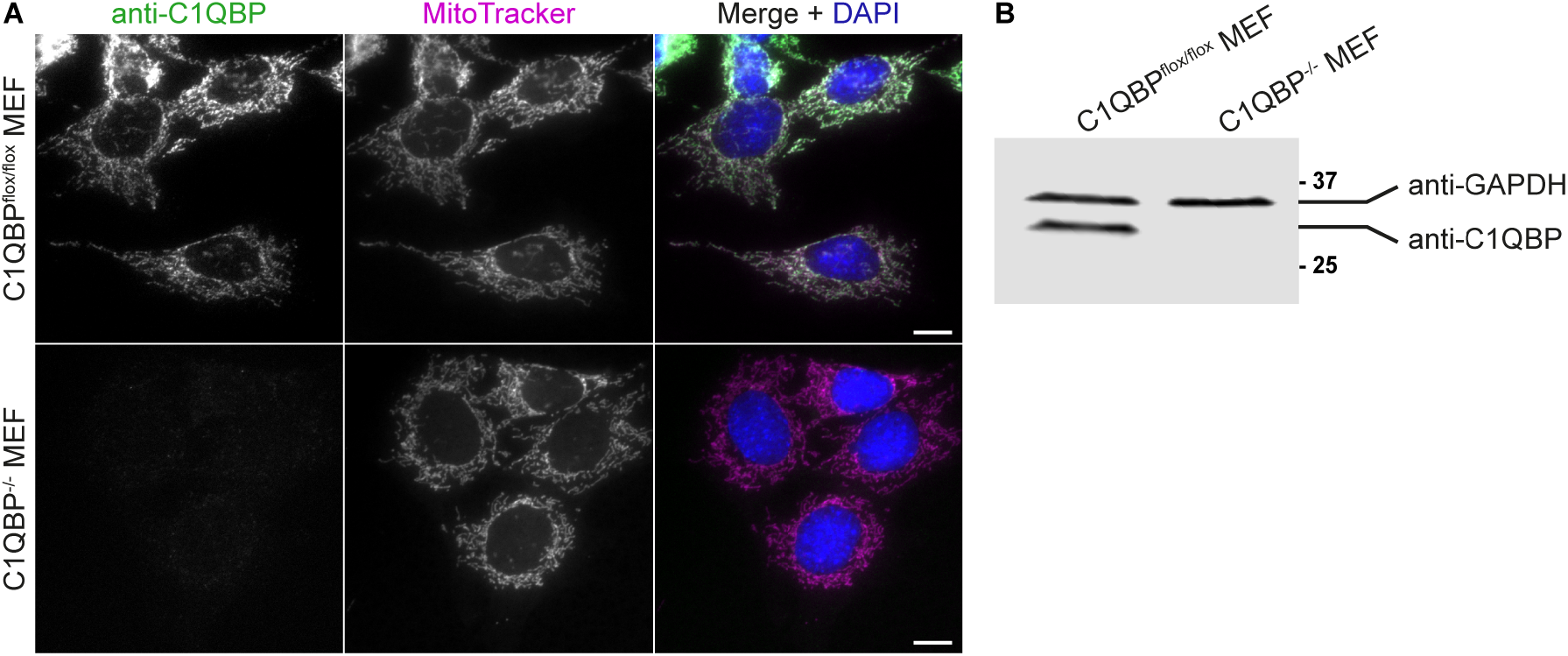
Verification of C1QBP^flox/flox^ and C1QBP^-/-^ MEFs. **A.** Immunofluorescence localisation of C1QBP and verification of knockout in C1QBP^-/-^ MEFs. Scale bar = 10 μm. **B.** Western blot verification of C1QBP knockout in C1QBP^-/-^ MEFs.

**Figure S8:**
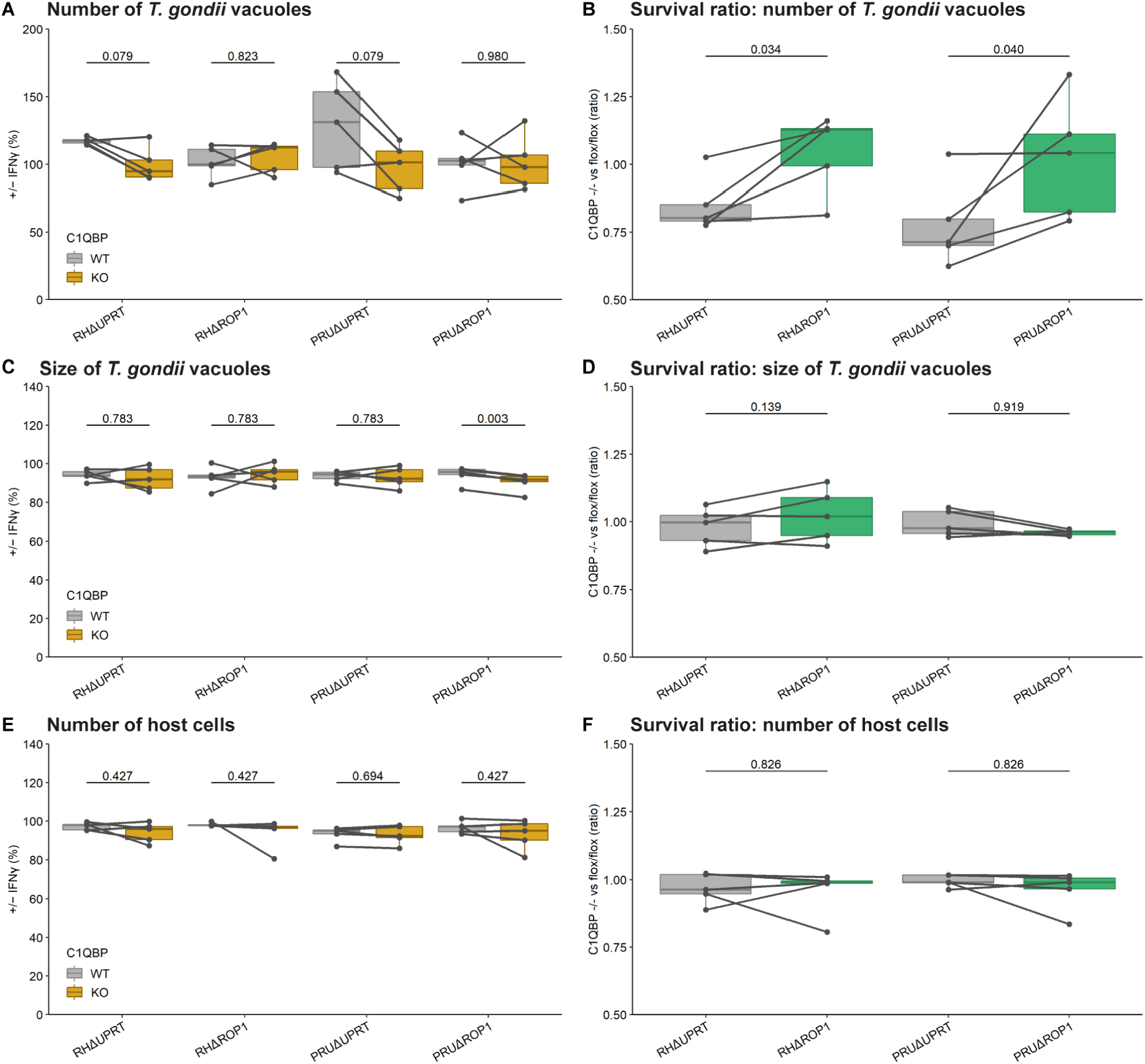
C1QBP knockout affects vacuole number, but not vacuole size or host cell death. **A, C, D.** IFNγ-dependent growth restriction of *T. gondii* in C1QBP^flox/flox^ (WT) and C1QBP^-/-^ (KO) immortalised MEFs. MEFs were stimulated with IFNγ for 24 h, infected with *T. gondii* cell lines for a further 24 h, and parasite growth quantified by automated fluorescence imaging and analysis. Parasite growth in IFNγ-stimulated BMDMs is shown as a percentage of that in unstimulated BMDMs in terms of **A** number of *T. gondii* vacuoles, **C** vacuole size, and **E** number of host cells. p-values were calculated by paired two-sided *t-*test with Benjamini-Hochberg adjustment. **B, D, F.** Ratio of the parasite survival in C1QBP^-/-^ versus C1QBP^flox/flox^ MEFs in terms of **B** number of *T. gondii* vacuoles +/− IFNγ, **D** vacuole size +/− IFNγ, and **F** number of host cells +/− IFNγ. p-values were calculated by paired two-sided *t-*test with Benjamini-Hochberg adjustment.

## SUPPLEMENTARY DATA

**Supplementary Data 1: CRISPR knockout screen results.**

**A.** Raw protospacer sequencing read counts.

**B.** Normalised protospacer sequencing read counts.

**C.** Protospacer L2FCs.

**D.** Gene L2FCs, p-values, and DISCO scores.

**E.** Comparison of L2FCs to (Young et al. 2019) and (Yifan Wang et al. 2020).

**Supplementary Data 2: BMDM IFNγ restriction assay results.**

**A.** *T. gondii* cell number, vacuole number, mean vacuole size and host cell number per well.

**B.** Median *T. gondii* cell number, vacuole number, mean vacuole size, host cell number and survival percentage +/− IFNγ per strain per replicate.

**Supplementary Data 3: THP-1 IFNγ restriction assay results.**

**A.** *T. gondii* cell number, vacuole number, mean vacuole size and host cell number per well.

**B.** Median *T. gondii* cell number, vacuole number, mean vacuole size, host cell number and survival percentage +/− IFNγ per strain per replicate.

**Supplementary Data 4: Co-immunoprecipitation mass spectrometry results.**

**Supplementary Data 5: MEF FNγ restriction assay results.**

**A.** *T. gondii* cell number, vacuole number, mean vacuole size and host cell number per well.

**B.** Median *T. gondii* cell number, vacuole number, mean vacuole size, host cell number and survival percentage +/− IFNγ per strain per replicate.

**C.** Ratio of survival in C1QBP^-/-^ vs. C1QBP^flox/flox^ MEFs.

**Supplementary Data 6: Primers sequences used in this work.**

**Supplementary Data 7: Opera Phenix image acquisition parameters and Harmony analysis sequence.**

**Supplementary Data 8: Microwave program used for TEM sample preparation.**

